# Multimodal imaging reveals no evidence for magnetite-based magnetoreceptors in the mole-rat eye

**DOI:** 10.64898/2026.04.02.715577

**Authors:** Leif Moritz, Kamalika Nath, Ella P. Walsh, Anna Sternberg, Elisabeth Becher, Adrian Lange, Gerald Falkenberg, Dennis Brückner, Clemens Diwoky, Kristian Bredies, Malte Brammerloh, Daryl Howard, David J. Paterson, Kadda Medjoubi, Stephan Irsen, Sybille Wolf-Kümmeth, Li Zhang, Martha M. M. Daniel, David A. Simpson, Sabine Begall, E. Pascal Malkemper

## Abstract

Magnetoreception, the ability to perceive the geomagnetic field, is widespread across animals. The underlying sensory mechanism remains elusive, but a long-standing hypothesis proposes single-domain magnetite linked to mechanosensitive ion channels. The Ansell’s mole-rat (*Fukomys anselli*) is a subterranean rodent with a magnetic sense, and published behavioral and histological data are consistent with magnetite-based magnetoreceptors in the cornea or retina. Here, we systematically screened for magnetite in the mole-rat eye, combining iron detection via enhanced Prussian blue staining and synchrotron X-ray fluorescence microscopy (XFM) with magnetic detection via MRI quantitative susceptibility mapping (MRI-QSM) and quantum-diamond microscopy (QDM). This revealed only a few iron particles in the retina and cornea, which predominantly overlapped with titanium or chromium, indicating a non-biogenic origin. XFM showed iron-enriched lines in the cornea, but these did not show ferrimagnetic signals. Focusing on other ocular tissues, MRI-QSM revealed the highest susceptibility in the ciliary body, where iron-rich pigmented cells were identified. A TEM-screen, however, failed to detect single-domain magnetite particles in these cells. We conclude that our high-sensitivity multimodal screen provides no evidence for magnetite-based magnetoreceptors in the mole-rat eye, suggesting that mole-rat magnetoreceptors either do not reside in the eye or are based on different physical principles.

## INTRODUCTION

Magnetoreception, the ability to sense the Earth’s magnetic field, is known from many organisms, including birds, bats, sea turtles, migratory fish, and subterranean mole-rats, which use it for navigation and orientation (Wiltschko and Wiltschko, 1972; Kimchi, Etienne and Terkel, 2004; Holland *et al*., 2006; Lohmann, Putman and Lohmann, 2012; Putman *et al*., 2013). Despite the widely accepted existence of magnetoreception, the receptor cells and mechanisms by which animals detect the magnetic field are still unknown (Nordmann, Hochstoeger and Keays, 2017). Several hypotheses have been proposed, three of which are currently supported by experimental evidence from different organisms. The electromagnetic induction hypothesis states that animals sense magnetic fields indirectly by detecting electric fields generated when they move through the Earth’s static magnetic field. As this requires a conductive loop and sensitive electroreceptors, it was, for a long time, only considered plausible for marine organisms (Jungerman and Rosenblum, 1980), but recent experimental evidence supports the idea in terrestrial animals (Nimpf *et al*., 2019; Nordmann *et al*., 2025). The radical pair hypothesis states that organisms possess quantum sensors in which the Earth’s magnetic field has an effect on chemical intermediates induced by photoexcitation (Hore and Mouritsen, 2016). This mechanism has gained considerable support in birds, with cryptochrome 4 as a candidate sensor protein, although the exact signal transduction mechanism has not yet been identified (Hochstoeger *et al*., 2020; Xu *et al*., 2021). The discovery of magnetotactic bacteria (MTB), which harbor a chain of single-domain (sd) magnetite (Fe_3_O_4_), turning the cells into miniature compass needles (Blakemore, 1975), gave rise to the third idea, which states that similar magnetite crystals, coupled to membranes or ion channels, could turn sensory cells in higher organisms into magnetoreceptors. The magnetite hypothesis of magnetoreception (Kirschvink and Gould, 1981; Shaw *et al*., 2015) has received some support from experiments in fishes, birds, bats, and mole-rats (Marhold, Wiltschko and Burda, 1997; Holland *et al*., 2008; Wiltschko *et al*., 2009; Naisbett-Jones *et al*., 2020). Furthermore, a hybrid sensor was proposed for night-migratory songbirds, in which magnetite nanoparticles might enhance the geomagnetic field effect on radical pairs by up to 100-fold (Hore, 2026). Regardless of the underlying mechanism, the prospect that magnetosensory cells exist in mammals is exciting because their molecular characterization could lead to a major breakthrough in magnetogenetics, i.e. the controlled activation of cells by magnetic fields (Nimpf and Keays, 2017).

Among mammals, magnetic sensing has been extensively investigated in Ansell’s mole-rat *Fukomys anselli* (Burda, Zima, Scharff, Macholán & Kawalika, 1999), a subterranean rodent from Zambia. Their ability to perceive the geomagnetic field, which they might use for orientation underground and as a heading indicator while digging (Malewski *et al*., 2018), was demonstrated by nest-building assays, in which the animals show a preference to nest in the magnetic southeast of a circular arena (Burda *et al*., 1990; Marhold, Wiltschko and Burda, 1997) (there referred to as *Cryptomys hottentotus*). Such experiments revealed that the magnetic sense of *F. anselli* functions in total darkness, is sensitive to the polarity of the magnetic field, and is affected by a short magnetic pulse but not by weak oscillating radiofrequency fields (Marhold *et al*., 1997; Marhold, Wiltschko and Burda, 1997; Thalau *et al*., 2006). These findings are in agreement with magnetoreceptors based on sd magnetite, similar to the nanocrystals found in MTBs (Blakemore, 1975; Frankel, 2009; Shaw *et al*., 2015). It is hypothesized that these crystals are coupled to the membrane of a mechanosensitive cell, where rotation or translation in a magnetic field would lead to their deflection and the opening of ion-channels (Fig. 1A). To act as a torque transducer in the weak magnetic field of the Earth (∼25-65 µT), magnetite has to be present in the form of single-domain particles, which have a size around 50-60 nm (Fig. 1A), or as collectives of smaller (20-50 nm) superparamagnetic crystals (Winklhofer and Kirschvink, 2010; Shaw *et al*., 2015). Sensory cells with such intracellular crystals have not been consistently identified in any vertebrate so far (Mouritsen, 2012). Considering the minute size (50 nm) of the potential magnetic particles and the relatively large size of *F. anselli*, with a body length of up to 13.5 cm (Begall, Burda and Caspar, 2021), screening the entire animal for magnetite is infeasible. Importantly, nest-building experiments also provided some clues on the potential location of the receptors. The directional preference in the nest-building assay was abolished when the eyes were surgically removed (Caspar *et al*., 2020) or when the cornea was locally anesthetized with lidocaine, with the latter treatment not affecting the detection of light (Wegner, Begall and Burda, 2006). Furthermore, ferric inclusions were detected in sections of the cornea of one mole-rat via Prussian blue staining, and the authors speculated that these might represent magnetoreceptive cells that contain magnetite (Wegner, Begall and Burda, 2006; Moritz *et al*., 2007). Interestingly, in bats, where behavioral data are consistent with a magnetite-based mechanism of magnetoreception (Holland *et al*., 2008), anesthesia of the cornea also affected orientation behavior (Lindecke *et al*., 2021). Moreover, in a transmission electron microscopy (TEM) study of the mole-rat eye, crystalloid bodies resembling magnetosomes of MTBs were reported in the outer nuclear layer (ONL) of the retina, but their elemental composition and diffractive or magnetic properties have not been investigated (Cernuda-Cernuda *et al*., 2003). Consistent with a magnetosensory function of the eye of mole-rats, a c-fos study reported neurons responsive to magnetic stimuli in the midbrain superior colliculus, which receives sensory input from the cornea and retina (Němec *et al*., 2001; Němec, Burda and Peichl, 2004).

**Figure 1:**
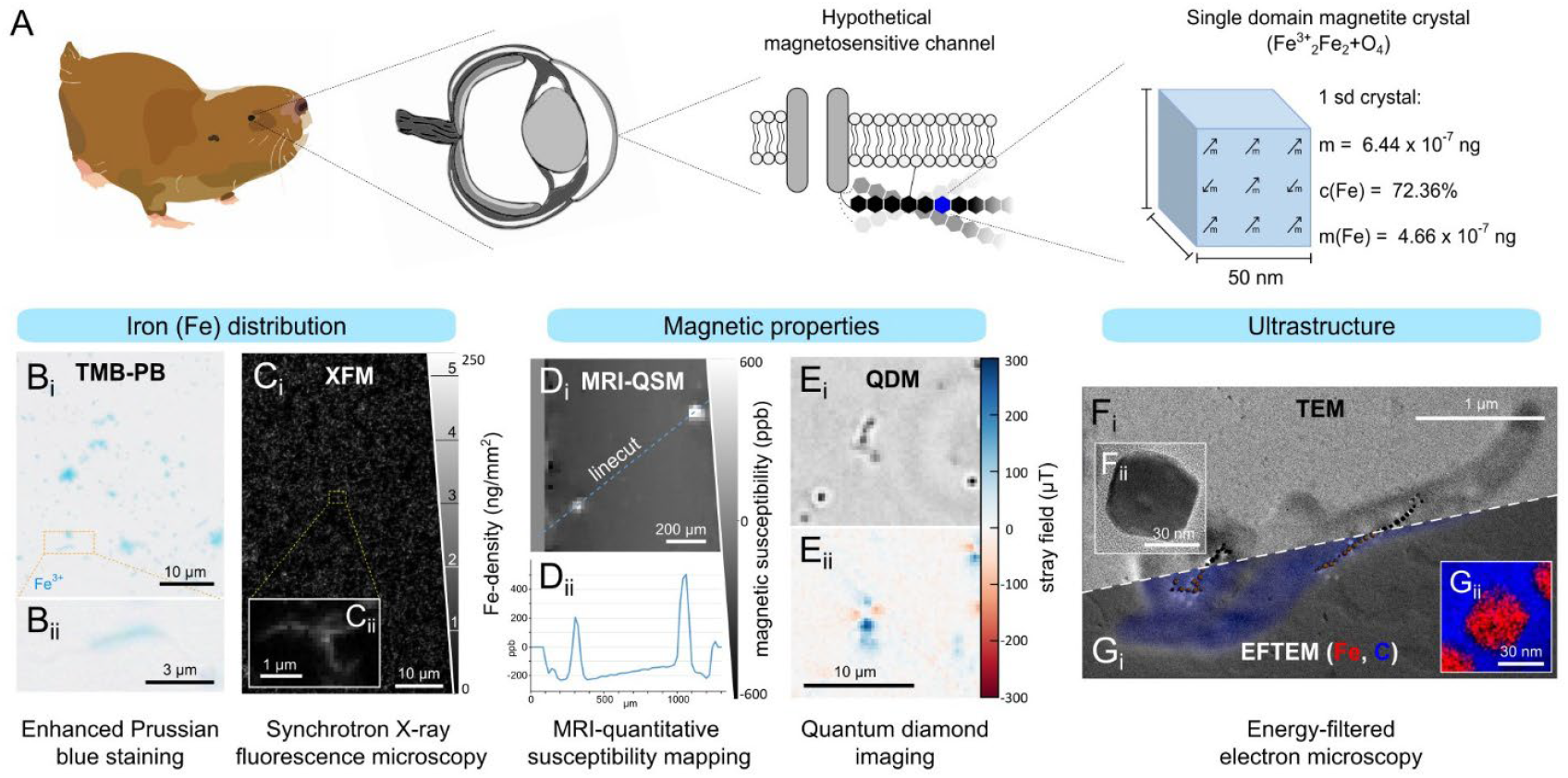
A multimodal imaging pipeline that enables magnetite detection across scales. **A** Hypothetical mechanosensitive magnetic-particle-based magnetoreceptor suspected in the eye of the Ansell’s mole-rat (*Fukomys anselli*) and the dimensions and theoretical iron (Fe) content of single-domain magnetite (Fe_3_O_4_) as found in magnetotactic bacteria (MTBs, not drawn to scale). **B – G** MTBs (*Paramagnetospirillum magnetotacticum*) served as a positive control to test the screening methods. **B** Positive TMB-enhanced Prussian blue staining of MTBs. **B**_**i**_ Overview image of stained MTBs. Note that some cultured MTBs turned ovoid and not all show the typical spirillum shape. **B**_**ii**_ Individual stained MTB. **C** Synchrotron X-ray fluorescence microscopy (XFM) gives quantitative density maps of Fe and other elements. **C**_**i**_ Overview of Fe-density in MTBs. Lines on the density-scale mark Fe densities expected for the corresponding number of single-domain (sd) magnetite crystals. **C**_**ii**_ Fe-density map of an individual MTB. **D** MRI-quantitative susceptibility mapping (MRI-QSM) detects MTB accumulations. **D**_**i**_ Single plane of MRI-QSM data with magnetic signals from MTBs. **D**_**ii**_ Linecut as indicated in B_i_ showing voxel intensity (ppb = parts per billion). **E** Quantum diamond microscopy (QDM) visualizes the magnetic dipole signals of individual MTBs. **E**_**i**_ Fluorescence image of MTB. **E**_**ii**_ Magnetic signal of the same MTB. **F** Transmission electron microscopy (TEM) allows the identification (F_i_) and structural analysis (F_ii_) of sd magnetite from MTBs. **G** Energy-filtered-TEM (EFTEM) shows elemental distribution within MTBs (G_i_) at high magnification (G_ii_).

The predominant method to screen for iron and potential magnetite in animal tissues in the context of magnetoreception is the Prussian blue (PB) staining (Perls, 1867), in which ferrous ferrocyanide reacts with ferric iron (Fe^3+^) in an acidic environment to form a bright blue ferric ferrocyanide complex (Fleissner *et al*., 2003, 2007; Falkenberg *et al*., 2010; Edelman *et al*., 2015; Shaw *et al*., 2018; Sonoda *et al*., 2025). The method has, however, been suggested to be unsuitable for finding sd magnetite crystals within cells, as it failed to yield positive results for magnetotactic bacteria (MTBs) (Curdt *et al*., 2022) and can lead to false positive results due to nonspecific staining of environmental contamination or haemosiderin deposits in macrophages (Treiber *et al*., 2012; Edelman *et al*., 2015). Furthermore, although Prussian blue staining can be used for coarse, rapid screening of iron in tissues, it does not provide information on the origin and magnetic properties of this iron, which are essential for characterizing magnetite-based magnetoreceptors.

Here, we employed a multimodal imaging pipeline to detect and characterize potential magnetite-based magnetoreceptors in animal tissues (Fig. 1). We developed an enhanced Prussian blue protocol that stains magnetite in MTBs blue (Fig. 1B) and complemented it with other imaging methods to map iron distribution and magnetic properties. Synchrotron X-ray fluorescence microscopy (XFM) gives detailed quantitative maps of iron (Fig. 1C) and other elements in tissue sections, while MRI-quantitative susceptibility mapping (MRI-QSM; Fig. 1D) and quantum diamond microscopy (QDM; Fig. 1E) probe the magnetic properties of tissues at different scales. Finally, TEM allows detailed examination of nanoparticles (Fig. 1F) and their elemental characterization, using energy filtering (EFTEM; Fig. 1G). We employed the pipeline on the eye of the mole-rat *Fukomys anselli*, focusing on the cornea and retina, which have been suggested to be involved in magnetoreception (Cernuda-Cernuda *et al*., 2003; Wegner, Begall and Burda, 2006; Caspar *et al*., 2020). We find no evidence of magnetite in the cornea or retina, or other parts of the eye.

## RESULTS

We undertook a multimodal screen for magnetite magnetoreceptors in the mole-rat eye. We first validated every method by confirming its ability to detect sd magnetite chains (magnetosomes) in the magnetotactic bacterium *Paramagnetospirillum magnetotacticum* (MTB; Fig. 1). We then applied this approach to screen for magnetite-based magnetoreceptors in the mole-rat eye by mapping iron distribution and magnetic properties across multiple scales.

### Prussian blue staining

We first aimed to replicate previous findings of abundant iron clusters in the epithelium of the cornea of mole-rat eyes (Wegner, Begall and Burda, 2006). Using the same standard Prussian blue (PB) iron-staining protocol, we manually screened sections of 12 corneas from 10 animals at regular intervals (10 µm thickness; 612 sections in total). We found very few Prussian blue-positive particles (40 ± 47 particles/cornea; 1 ± 1 particles/section, mean ± standard deviation (SD)) and high variance between individuals (Fig. S1A). Particles appeared markedly different from the large, regular, and abundant particles reported in the epithelium of a single cornea section by Wegner et al. (2006). We found particles across all corneal layers (n = 10 animals, 12 eyes, Fig. S1A & B), and a reconstruction of the particle positions from serial sections revealed no regular pattern in their distribution across the cornea (n = 4 animals, Fig. S1B). In summary, using similar methods, we were unable to replicate the histological findings reported by Wegner et al. (2006).

Recently, doubts were raised on the use of PB staining as a screening tool for animal magnetoreceptors, because it does not visualize the magnetite chains of magnetotactic bacteria (Curdt *et al*., 2022). We observed the same lack of staining with the classical PB protocol (Fig. S2A). To amend this problem, we developed an amplified PB protocol. Based on a previously described amplification of PB with diaminobenzidine (DAB), which creates a brown stain (Nguyen-Legros *et al*., 1980; Perl and Good, 1992; Moos and Møllgård, 1993), we reasoned that other benzidine-based chromophores might work and found 3,3’,5,5’-tetramethylbenzidine (TMB) to react with PB. This produced an intense blue stain, stronger than the original PB reaction product, enabling us to visualize intracellular iron in individual MTBs with light microscopy (Figs 1E, S2B). Electron microscopy of stained MTBs revealed that the crystals were partially dissolved and that the iron spread within cells after treatment with 1% HCL, and after classical PB staining (Fig. S2 D&G), as well as after the TMB-enhanced PB-staining (Fig. S2E&H). This suggests that, although the PB reaction occurs under the classical protocol, the resulting precipitate is insufficient for detection by light microscopy without signal enhancement. The intense blue of TMB enhancement is advantageous over the brown of DAB when screening for iron in pigmented tissues such as the ciliary body and pigment epithelium of the eye (Figs 2A, S1F).

**Figure 2:**
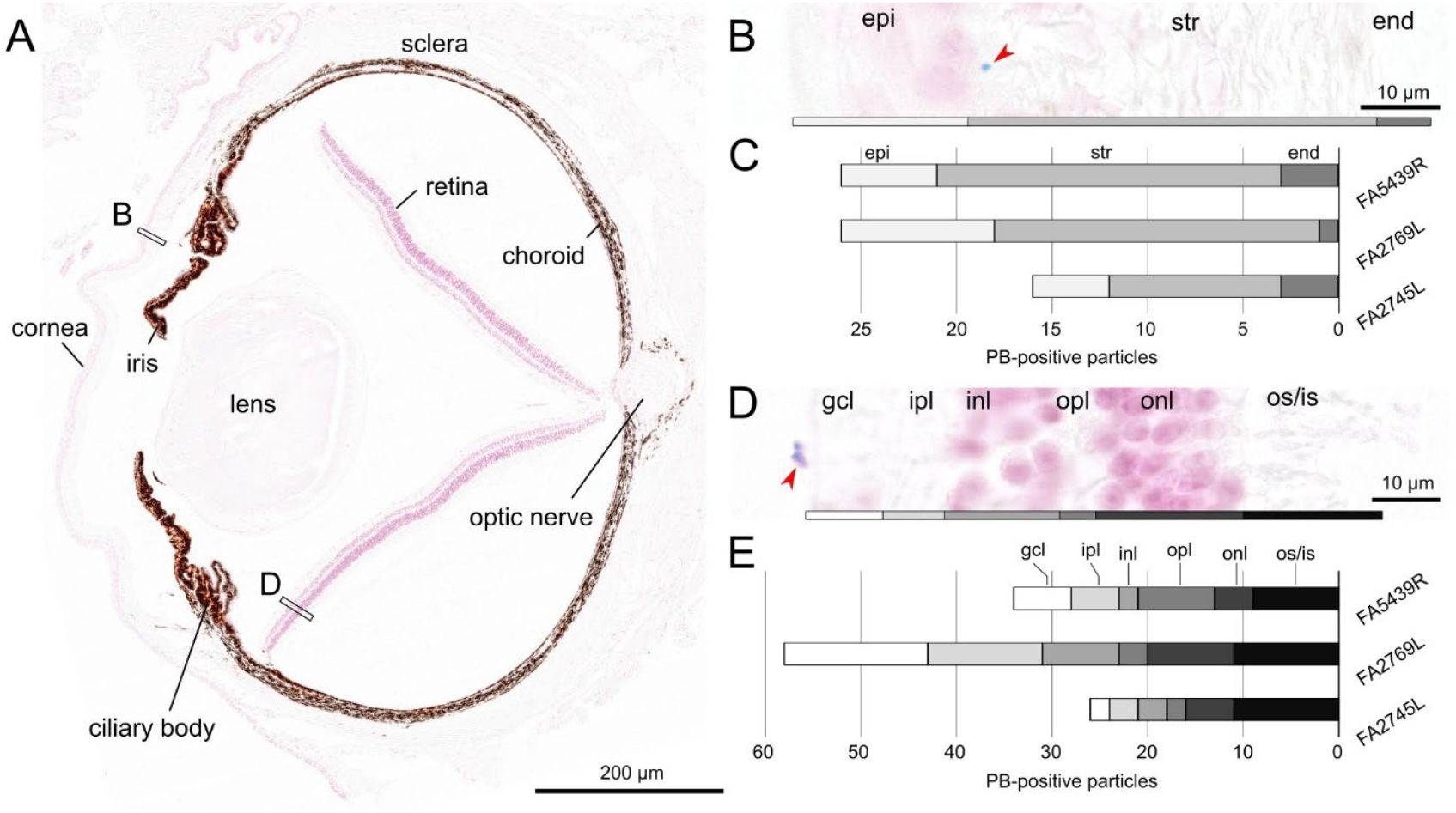
Enhanced Prussian-blue (TMB-PB) staining in serial sections of three mole-rat (*Fukomys anselli*) eyes reveals rare iron particles in the cornea and retina randomly distributed across all layers. A total of 960 sections of 3 animals have been screened. **A** Section of a mole-rat eye stained with TMB-PB and counterstained with nuclear fast red. The V-shaped detached retina is an artifact resulting from the paraffin embedding. **B** Detail of the cornea (box B indicated in A) with exemplary TMB-PB-positive particle in the epithelium indicated by red arrow, shown at the 100x magnification at which the screen was performed. **C** Quantification and location of TMB-PB positive particles in the cornea of three animals (Number of sections: FA5439R = 365; FA2769L = 291; FA2745L = 304). **D** Detail of the retina (box D indicated in A) with exemplary TMB-PB-positive particle in the ganglion cell layer. **E** Quantification and location of TMB-PB-positive particles in the retina of three animals (Number of sections: as for C). Abbreviations: end = endothelium, epi = epithelium, gcl = ganglion cell layer, inl = inner nuclear layer, ipl = inner plexiform layer, onl = outer nuclear layer, opl = outer plexiform layer, os/is = outer/inner segments, str = stroma.

Using the TMB-enhanced PB protocol, we manually screened all serial sections (Figs 2A, S3) of three mole-rat eyes from three individuals for iron at high magnification (5 µm thickness, n = 3 eyes, 960 sections in total). We detected 23 ± 5 Prussian blue positive particles (mean ± SD, n = 3 eyes) in the cornea (epithelium = 6 ± 2; stroma = 15 ± 4; endothelium = 2 ± 1; Figs 2B&C, S3), and 39 ± 17 particles in the retina (OS/IS = 10 ± 1; ONL = 6 ± 3; OPL 4 ± 3; INL = 4 ± 3; IPL = 7 ± 5; GCL = 8 ± 7; Fig. 2D&E). Again, the distribution across layers appeared random in the cornea and retina, and no consistent pattern could be observed across individuals (Fig. 2C&E). The appearance and size of the particles were highly variable (Fig. S3). We also screened other parts of the eyes and found 5 ± 5 particles in the ciliary body, 0 ± 1 in the iris, 7 ± 6 in the lens, 17 ± 7 in RPE, and 141 ± 29 in the sclera (Fig. S1C–F).

In sum, using both standard and enhanced PB staining, we did not find the previously reported abundant iron rich clusters in the mole-rat cornea (Wegner, Begall and Burda, 2006), and only a few particles in the retina and several other ocular tissues. We next asked whether these particles were of biological origin or contamination from the laboratory environment (Edelman *et al*., 2015; Malkemper *et al*., 2019). To address this, we investigated their elemental composition quantitatively using synchrotron X-ray fluorescence microscopy (XFM).

### Synchrotron X-ray fluorescence microscopy (XFM)

We performed elemental screening of mole-rat eye sections using XFM at three synchrotrons: ANSTO (Australia), DESY (Germany), and Soleil (France) (Fig. 3). To do this, we mounted 2 µm thick epon sections of mole-rat eyes on silicon nitride windows and scanned them at the XFM beamlines. These measurements revealed the highest spatial Fe-density in the ciliary body (38.04 ± 11.18 ng/cm^2^, n = 4 animals, 35 sections), iris (31.51 ± 17.74 ng/cm^2^, n = 4 animals, 11 sections), and the choroid (20.30 ± 2.16 ng/cm^2^, n = 4 animals, 36 sections). Considerably lower average densities were found in the sclera (4.43 ± 2.98 ng/cm^2^, n = 4 animals, 36 sections) and cornea (4.26 ± 2.56 ng/cm^2^, n = 4 animals, 19 sections). The lens (2.71 ± 4.17 ng/cm^2^, n = 3 animals, 24 sections) and retina (1.11 ± 0.63 ng/cm^2^, n = 3 animals, 14 sections) had the lowest Fe-densities (Tab. S1).

**Figure 3:**
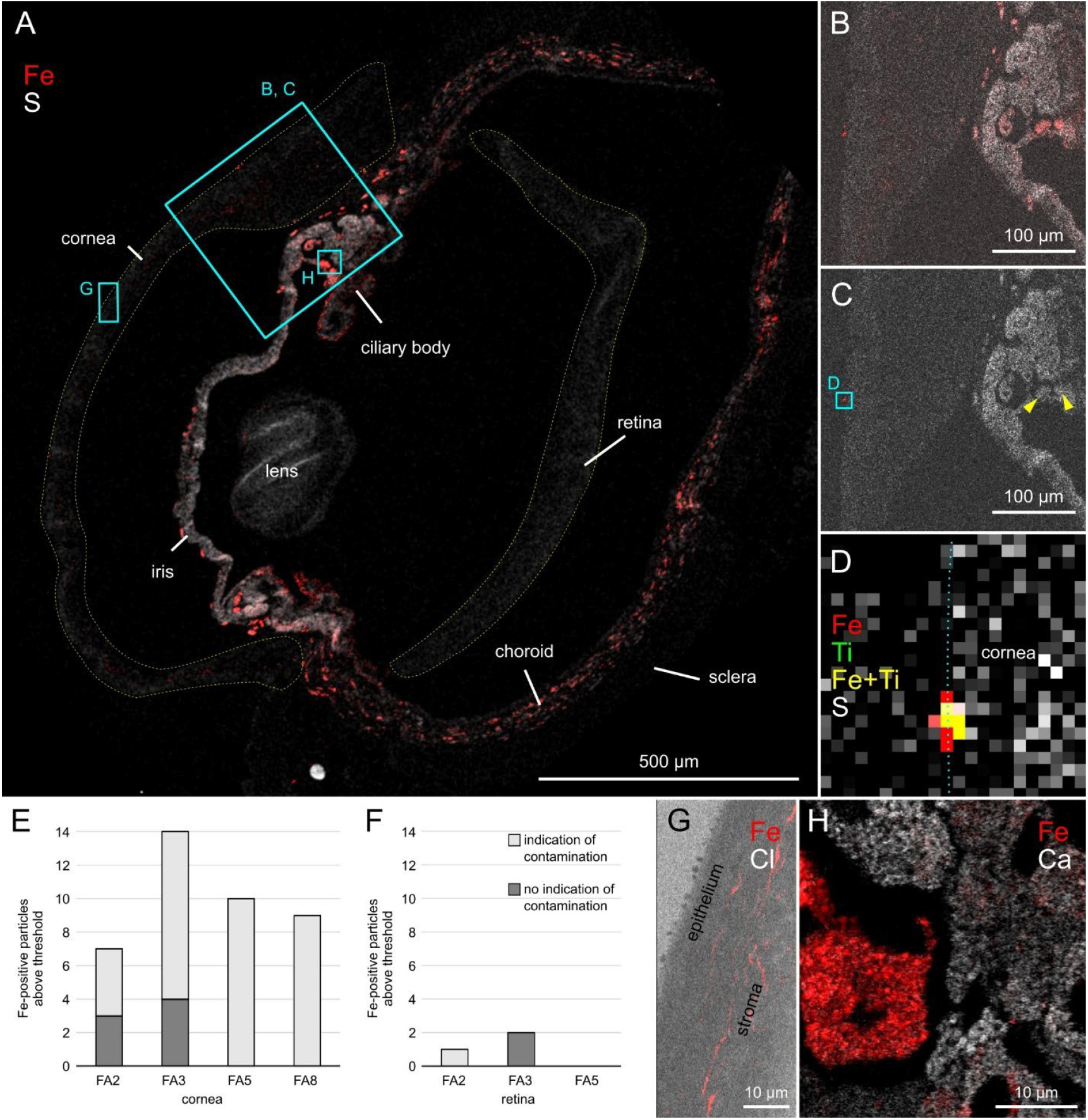
Synchrotron X-ray fluorescence microscopy of mole-rat (*Fukomys anselli*) eye sections revealed contamination in the cornea and retina and iron-enriched cells in the corneal stroma and ciliary body. **A** Overview of an eye section (ID: 9FA5, section 14) showing Fe-density (red) on Sulfur(S)-background. Dotted lines indicate the cornea and retina. The detached retina is a histological artifact. Data obtained at ANSTO. **B** Detail of A, showing Fe-density in the cornea and ciliary body. **C** Same area as in B, but only highlighting pixels with Fe-content above 20 sd magnetite crystals (50 nm cubes). Yellow arrows indicate Fe-rich particles in the ciliary body. **D** Overlap of Fe- and Ti-in a Fe-rich particle in the cornea (area indicated in C), blue dotted line indicates surface of cornea, 1 pixel = 1 µm. **E & F** Quantification of particles with Fe-density ≥ Fe expected in a pixel with 20 sd magnetite crystals, with and without signs of contamination (Ti or Cr) in 4 individuals (ID on x-axis). **E** Corneas of 4 animals (n = 19 sections; FA2 = 6; FA3 = 4, FA5 = 4, FA8 = 5). **F** Retinas of 3 animals (n = 14 sections; FA2 = 5; FA3 = 5; FA5 = 4). **G** Sub-threshold Fe-lines in the stroma of the cornea, which were present in all animals and did not co-localize with markers of contamination. **H** Fe-rich cells in the ciliary body on Calcium (Ca)-background, approximate position as indicated in A. High-resolution data obtained at the Soleil synchrotron Nanoscopium beamline, 1 pixel = 150 nm.

Next, following established methodology (Malkemper *et al*., 2019), we identified candidate magnetoreceptor structures by selecting pixels with an Fe-density above or equal to the density expected in a pixel containing 20 sd magnetite particles (total mass = 128.8 × 10^-7^ ng, Fe-mass = 93.2 × 10^-7^ ng, Fig. 1A-C), which would be sufficient to open mechanosensitive ion channels (Winklhofer and Kirschvink, 2010; Goychuk, 2015; Malkemper *et al*., 2019). Subsequently, we checked candidate pixels by quantitating titanium (Ti) and chromium (Cr), which are common in contamination from the laboratory (Fig. 3D).

Within the cornea, we identified in total 40 particles above the threshold (9±3 particles per animal, n = 4 animals, 19 sections) of which 33 (82.5%) were co-localizing with elevated Ti- and/or Cr-densities (> background-mean-density + 2 SD). Notably, in two animals, all particles co-localized with markers of contamination (Fig. 3E), and particles without contamination were predominantly found at the margin of the cornea to other tissues (mainly the Fe-rich ciliary body, Fig. S4D&E). In the retina, we detected only three particles above the threshold (1±1 per animal, n = 3 animals, 14 sections), of which one co-localized with elevated Cr-density (Fig. 3F). Of the remaining particles, one was located on the surface of the retina facing the vitreous body (Fig. S4E) and one was located underneath the retina, which detached from the RPE/choroid (Fig. S4F).

Based on these data, we concluded that the particles detected in the cornea and retina with PB staining were contaminants rather than magnetoreceptors. In other parts of the eye, we noted the highest fraction (36.8 ± 8.0%) of non-contaminated iron-rich regions in the ciliary body (427 particles, n = 4 animals, 35 sections; Fig. S4). Interestingly, high-resolution XFM imaging at the Soleil synchrotron revealed cells in the pigmented ciliary epithelium filled with iron-rich granules (Fig. 3H), typically surrounded by neighboring cells enriched in zinc (Zn).

Moreover, at a lower threshold for iron concentration, XFM revealed regular Fe lines in the corneal stroma (Fig. 3G), which were consistently observed across all samples. These lines were not associated with contamination but remained below the density expected for 20 sd magnetite particles (n = 4 animals, 19 sections). To investigate their magnetic properties, we next performed quantum diamond microscopy on the cornea.

### Quantum diamond microscopy (QDM)

To probe the magnetic properties of the cornea at high spatial resolution, we used quantum diamond microscopy (QDM). QDM leverages spin-dependent fluorescence of nitrogen vacancy (NV) centers in diamond to detect magnetic signals with high spatial resolution and at low applied magnetic fields (∼4 mT). Here, we used optically detected magnetic resonance (ODMR) to map the cornea, which, via XFM, consistently showed lines of sub-threshold iron densities in the stroma across all samples (Figs 4, S5).

**Figure 4:**
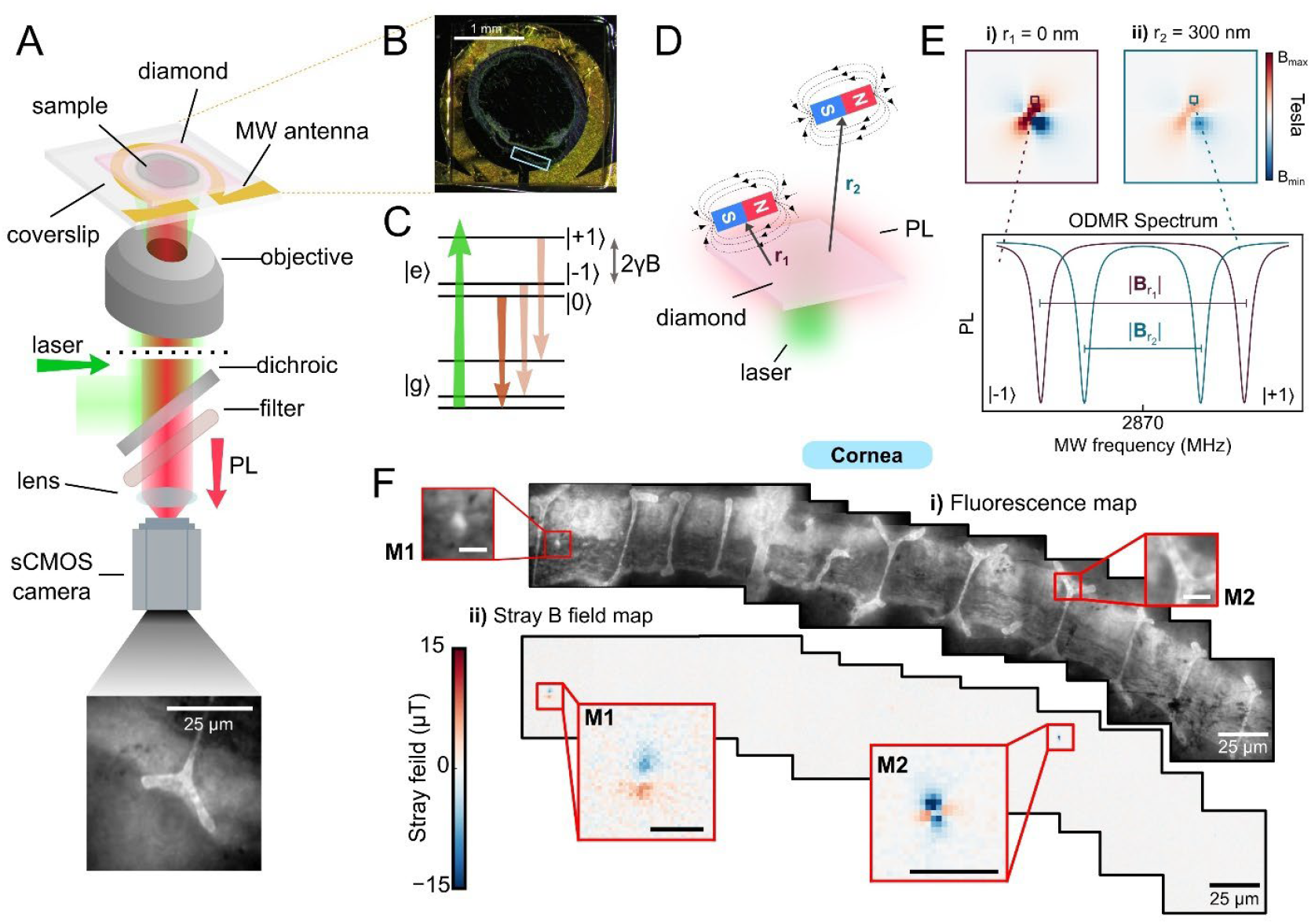
Quantum diamond microscopy revealed (exogenous) ferrimagnetic signals in the cornea but not in sub threshold iron-lines. **A** Experimental scheme of the Quantum Diamond Microscope. **B** 500 nm tissue slice on a diamond sample. **C** Energy levels of the nitrogen vacancy (NV) center in diamond used to map the local magnetic fields. **D** Simulated magnetic field strength from a dipole scaled as |B|∝r^3^. **E** Simulated ferromagnetic signal from sd magnetite with distance. **F** Optical fluorescence images and corresponding stray magnetic field maps of the cornea. M1 & M2 are the two magnetic signals found in the tissue at an applied field of ∼4 mT. Vertical bright lines are folds, where the sample is not attached to the diamond. Scales in details of M1 & M2 = 5 µm.

Photoluminescence (PL) maps revealed a total of two prominent magnetic signals in the cornea (1 animal, 2 sections, 25 FOVs; Figs 4F (M1 & M2), S4), but no signal corresponded in shape or position to the XFM-detected lines. The stray magnetic signals had varying dipole orientations and peak-to-peak strengths of ∼12 μT (M1) and ∼20 μT (M2) (Figs 4F, S5). Comparison with magnetite chains in magnetotactic bacteria (MTBs) under an applied field of 4 mT showed stronger stray fields (∼28 μT), as expected, because the magnetite is directly at the diamond surface (Fig. S6).

The fact that the magnetic dipoles in the cornea were not aligned with the applied field (Figs 4, S5) indicates that the signals were not paramagnetic, but possess intrinsic ferro/ferrimagnetic moments. Comparison to the fluorescence images showed that M1 corresponded to a structure considerably larger than sd magnetite, while M2, the strongest signal (∼20 μT), was located on or beneath a fold in the section (Figs 4F, S5), indicating likely contamination. These observations confirm that QDM can reliably detect ferro/ferrimagnetic particles and demonstrate that the sub-threshold Fe-lines consistently observed in the corneal stroma are not particle-based magnetoreceptors. The cornea was thus excluded from further consideration, and we next focused on other regions of the eye.

### Magnetic resonance imaging-quantitative susceptibility mapping (MRI-QSM)

To identify additional potential magnetoreceptor sites in the mole-rat eye, we next examined the volume magnetic properties of ocular tissues using magnetic resonance imaging quantitative susceptibility mapping (MRI-QSM). Scans were acquired at an isotropic resolution of 20 µm (n = 7 eyes from 7 animals; Fig. 5A&B). While simulations following (Brammerloh *et al*., 2024) revealed that our scans had a lower detection limit of 55 sd magnetic particles in a voxel (see Supplemental Materials), it allowed for comparisons of the bulk susceptibility of different ocular tissues.

**Figure 5:**
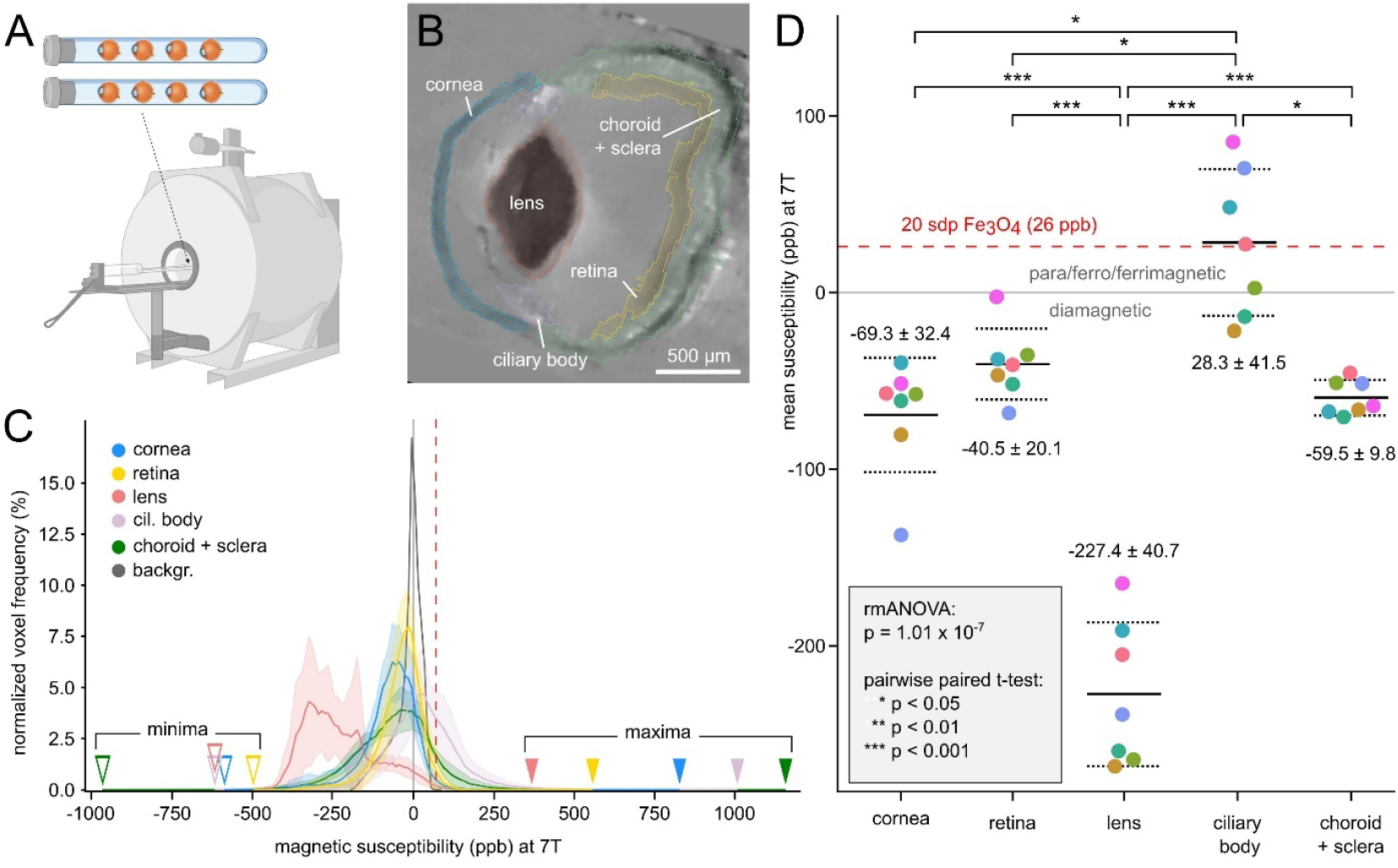
MRI-quantitative susceptibility mapping (MRI-QSM) of mole-rat eyes revealed the strongest magnetic signals in the ciliary body. **A** Eight eyes were measured simultaneously at 7 T yielding volume susceptibility maps of 20 µm isotropic resolution. One eye was excluded due to artifacts (Figure created in BioRender. Moritz, L. (2026) https://BioRender.com/r8md8a5). **B** Virtual section of MRI-QSM measurement of an eye with the areas analyzed indicated (colors correspond to C). **C** Histogram of the mean ± SD (n = 7) voxel frequency binned to 10 ppb magnetic susceptibility intervals in different tissue in the 7 eyes. **D** Mean susceptibility of the cornea, retina, lens and ciliary body across seven individual eyes. Colors indicate individuals. The magnetic signals from the ciliary body were significantly larger than in the other investigated tissues. The dashed red lines indicate the 26 ppb level expected for 20 single-domain (sd) magnetite crystals present in a voxel (see supplement). rmANOVA output was Greenhouse-Geisser-corrected for sphericity, t-test output was Bonferroni-corrected for multiple testing, non-significant comparisons are not shown.

Qualitatively, the distribution of the magnetic signals in the eyes matched the iron distribution detected with XFM, with low susceptibility in retina, cornea and lens, and higher values in the ciliary body (Fig. 5C-D). The susceptibility of different ocular tissues differed significantly (rmANOVA, n = 7, degrees of freedom = 4, F = 88.24, p = 1.01 x10^-7^ after Greenhouse–Geisser correction for violation of sphericity (ε = 0.485)). On average, the cornea (-69.3 ± 32.4 ppb), retina (-40.5 ± 20.1 ppb), lens (-227.4 ± 40.7), and choroid + sclera (-59.5 ± 9.8) were diamagnetic (< 0 ppb). On the contrary, the ciliary body (28.3 ± 41.5 ppb) showed positive (> 0 ppb) and significantly higher susceptibility values (post-hoc pairwise t-test: p < 0.05 after Bonferroni correction of p-values for multiple testing (6) between all tissue pairs except cornea-retina, cornea-choroid+sclera, retina-choroid+sclera (p > 0.5)), indicative of para- or ferro/ferrimagnetic content (Fig. 5D). However, these magnetic measurements are performed under an extremely high magnetic field (∼ 7 T) which is strong enough to magnetize a variety of magnetic materials such as magnetite, ferritin and ferrihydrite that exist within tissue, resulting in a combined magnetic response for entire regions of tissue. Furthermore, as the susceptibility is summed up over a voxel volume of 8,000 µm^3^, signals of magnetite might be masked by surrounding structures with a generally low (or negative) susceptibility, and it cannot be discerned whether high susceptibilities are due to highly ferrimagnetic but minute magnetic particles like single-domain magnetite, or larger structures with lower paramagnetic signal.

### Energy-Filtered Transmission Electron Microscopy (EFTEM)

To ascertain whether the high iron content (XFM) and magnetic signals (MRI-QSM) in the ciliary body originated from chains of sd magnetite crystals, we performed high-resolution imaging of the iron-enriched cells in the ciliary body using transmission electron microscopy. Using a magnification and settings that allowed us to resolve sd magnetic crystals in MTBs (8000x, pixel size = 1.5 nm, Figs 6A-C, S2C&F), we screened 93 fields of view (each 6 × 6 µm) in two mesal sections of two animals via energy filtered TEM (EFTEM). Elemental mapping based on EFTEM confirmed that these cells were rich in iron, with the iron appearing evenly spread in electron-dense pigment granules (Fig. 6D-F). We did not find evidence for iron concentrated in sd magnetite crystals, as those found in MTBs (Fig. 6A-C).

**Figure 6:**
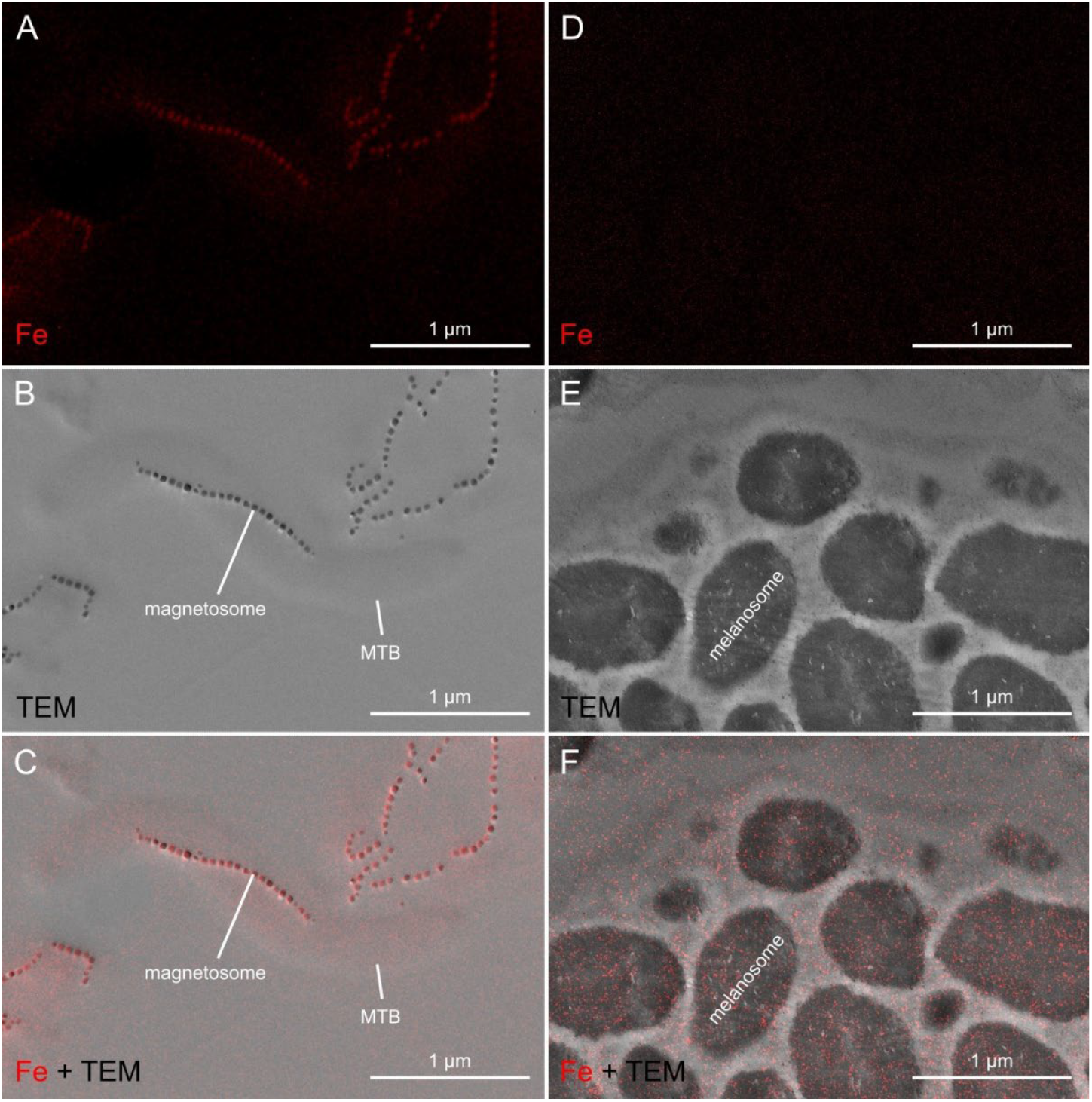
TEM and EFTEM screening of cells in the ciliary body revealed iron-rich melanosomes, but no indication of single-domain magnetite chains. **A – C** EFTEM image showing magnetotactic bacteria with iron concentrated in magnetosomes **A** Fe-jump ratio. **B** TEM image. **C** Overlay of Fe-jump ratio (red) and TEM image (gray). **D – F** EFTEM image taken with same settings as A-C, showing iron within cells with a diffuse iron distribution and no clear concentration in well-defined crystals as in magnetosomes **D** Fe-jump ratio. **E** TEM image. **F** Overlay of Fe-jump ratio (red) and TEM image (gray).

## DISCUSSION

In this study, we present a multimodal imaging approach to screen animal tissue for biogenic magnetite. We demonstrate that the methods can detect intracellular sd magnetite crystals, using magnetotactic bacteria (MTBs) as a positive control. Applying this approach to the mole-rat eye, we find no evidence of sd magnetite chains in the cornea, retina, or any other ocular tissue examined.

### Enhanced Prussian blue screening for the detection of sd magnetite

Prussian blue staining (Perls, 1867) is the predominant method to screen for iron in the context of magnetoreception (Fleissner *et al*., 2003, 2007; Falkenberg *et al*., 2010; Edelman *et al*., 2015; Shaw *et al*., 2018), but the sensitivity of the method to screen for iron in nervous tissue has been questioned (Perl and Good, 1992), as it failed to yield positive results for magnetotactic bacteria (MTBs) (Curdt *et al*., 2022) and led to false positive results due to contamination or macrophages (Treiber *et al*., 2012; Edelman *et al*., 2015). Contrary to these findings, a recent study demonstrated that MTBs, which penetrated into tumor cells, lead to positive intracellular Prussian blue staining, although this study used ovoid marine magnetotactic bacteria MO-1 and not spirillia (Wang *et al*., 2024). It is possible that the size, size ratio, or composition of magnetosomes in different bacterial species accounts for the reported differences in staining efficiency. We here confirm the results of Curdt et al. (2022), that classical Prussian blue staining does not lead to a visible blue precipitation in MTBs of the species *Paramagnetospirillum magnetotacticum*, but that a previously reported DAB-amplification (Nguyen-Legros *et al*., 1980; Perl and Good, 1992; Moos and Møllgård, 1993) leads to a brown stain. The here established TMB-amplification leads to blue intracellular precipitation after classical Prussian blue staining in these MTBs. That TMB enhancement yields a positive result in MTBs indicates that the classical Prussian blue reaction occurs, but the original staining product is insufficient to be detected with the light microscope. The advantage of the TMB enhancement is that the blue staining product is also clearly visible in (brown)-pigmented tissue, such as the eye.

### No evidence for magnetite-based magnetoreceptors in the cornea and retina

Behavioral studies on *F. anselli* are consistent with magnetite-based receptors (Marhold *et al*., 1997; Marhold, Wiltschko and Burda, 1997; Thalau *et al*., 2006), and support a role of the eyes in magnetoreception (Wegner, Begall and Burda, 2006; Caspar *et al*., 2020). Furthermore, previous studies reported inclusions with ferric iron in the cornea of *F. anselli* via classical Prussian blue staining (Wegner, Begall and Burda, 2006) or detected crystalloid bodies in the retina via TEM (Cernuda-Cernuda *et al*., 2003), speculating about their potential role in magnetoreception. Therefore, we expected to find sd magnetite as part of the putative magnetoreceptors in the cornea or retina of *Fukomys anselli*.

While our iron screen detected ferric iron (Fe^3+^; via enhanced Prussian blue-staining) in the cornea and retina of the Ansell’s mole-rat’s eye, it was not as abundant and ordered as previously reported for the corneal epithelium (Wegner, Begall and Burda, 2006), and it was distributed randomly across the cornea, which is inconsistent with a sensory structure. Furthermore, the detected particles mostly overlapped with titanium and/or chromium, indicating an external non-biogenic origin (contamination), or were located at tissue margins (e.g., on the surface of the cornea and retina, or adjacent to the ciliary body), suggesting that these are not native to the cornea and retina. Previous studies, using Prussian blue staining, demonstrated that accumulations of iron in the cornea (of humans), and especially in its epithelium, are not uncommon and can be related to aging or diseases (Jones and Reid, 1984; Loh, Hadziahmetovic and Dunaief, 2009; Wróblewska *et al*., 2024). Furthermore, environmental contamination (especially in substrate-digging animals like African mole-rats) might introduce exogenous iron into tissues, as proposed for the nasal epithelia of fish (Curdt *et al*., 2022). Finally, iron is sequestered by macrophages which are common across tissues (Treiber *et al*., 2012; Edelman *et al*., 2015; Sonoda *et al*., 2025). As iron is an abundant trace element in animal tissue, including the eye (Jones and Reid, 1984), and on its own is not indicative of sd magnetite crystals, we also took the magnetic properties of the tissue into account. As shown by MRI-QSM, the cornea and retina are overall diamagnetic and have a lower susceptibility than other iron-rich tissues, such as the ciliary body. However, as the sum of the susceptibility in an 8,000 µm^3^ voxel was measured, magnetite particles might be masked by diamagnetic surrounding tissue, leading to a diamagnetic signal despite their presence (false negatives), or weakly paramagnetic structures could add up, leading to high susceptibility despite their absence (false positives). This limitation was overcome by quantum diamond microscopy, which allows screening for magnetic properties at subcellular resolution and detected two particles with a ferro/ferrimagnetic signal in the cornea similar to that of MTBs. However, the position or appearance in the fluorescence image were indicative of other sources than biogenic sd magnetite crystals (e.g. environmental contaminations). Furthermore, subthreshold iron-lines consistently found via XFM in the cornea were found not to be ferro/ferrimagnetic. In summary, using a complementary multimodal approach that screens for iron and magnetic signatures across multiple resolutions and sensitivities, we find no evidence for sd magnetite as part of a magnetoreceptor in the mole-rat cornea or retina.

### Iron rich cells in the ciliary body

The large-scale XFM-screening and eye-wide MRI-QSM showed relatively high iron concentration and overall higher susceptibility in the ciliary body. However, the source is unlikely (biogenic) magnetite associated with a sensory structure, for several reasons. Firstly, Prussian blue staining revealed only few particles, which could be due to iron being ferrous (Fe^2+^) or tightly bound to proteins, as Prussian blue is only sensitive to ferric (Fe^3+^), non-heme iron (Meguro *et al*., 2007; Sonoda *et al*., 2025). Second, in contrast to the cornea and retina, the ciliary epithelium is highly vascularized and produces the aqueous humor, and it plays an active role in iron storage and transport (Ashok *et al*., 2018). Third, iron could stem from the strong pigmentation, as melanin is known to bind iron (Kaczara *et al*., 2012). Finally, iron particles have been reported in the ciliary body and sclera of humans, related to hemochromatosis (Roth and Foos, 1972; Jones and Reid, 1984), a disease that leads to an increased intake and accumulation of iron. Accumulations of excess iron in the liver (hepatic hemosiderosis), but no direct evidence for hemochromatosis, have been documented in captive naked mole-rats *Heterocephalus glaber* (Delaney, Imai and Buffenstein, 2021). In line with these arguments, our electron microscopic screening revealed no evidence of magnetite crystals in the ciliary body of *F. anselli*, indicating that these cells are unlikely to contribute to magnetoreception.

The iron-rich cells in the ciliary body are still noteworthy, since the blind mole rat *Nannospalax ehrenbergi*, another subterranean and magnetoreceptive rodent (Kimchi and Terkel, 2001; Kimchi, Etienne and Terkel, 2004) (therein as *Spalax ehrenbergi*), has been reported to have an hypertrophied iris-ciliary body complex despite otherwise degenerative features of the eye (Sanyal *et al*., 1990). Furthermore, the subterranean African mole-rat *Bathyergus suillus* shows a disproportionally large and highly pigmented ciliary body (Nikitina and Kidson, 2014). This could indicate that the ciliary body still plays an important physiological role, even though the eye does not respond to light stimuli (Sanyal *et al*., 1990).

## Conclusion

We present a multimodal screening approach for magnetite in animal tissues that combines methods sensitive to iron and magnetic properties and is applicable to a wide range of tissues and organisms, for example, to screen for the proposed hybrid magnetite-radical-pair sensor in the eyes of night-migratory songbirds (Hore, 2026). Using this comprehensive screening strategy, we found no evidence for magnetite-based magnetoreceptors in the eye of the mole-rat *Fukomys anselli*. It is possible that magnetite-based receptors are located in other tissues, and that the eyes are not involved in magnetoreception. Previous behavioral manipulations, such as eye removal (Caspar *et al*., 2020) and lidocaine anesthesia (Wegner, Begall and Burda, 2006) should be interpreted cautiously, as they may produce nonspecific behavioral side effects and could also affect surrounding tissues, adjacent nerves, and the central nervous system (Engels et al. 2018).

We conclude that mole-rat magnetoreceptors are either likely located in tissues outside the eye or are based on a mechanism other than magnetic particles, highlighting the need for further research and a re-evaluation of current models of magnetoreception in mammals. Although it is currently the best-supported hypothesis, magnetoreception in *F. anselli* may not be magnetite-based, and other mechanisms should be considered. An alternative mechanism would be electromagnetic induction, where an electric field is induced by movement through the static geomagnetic field. This mechanism does not require light and has recently been suggested to take place in the inner ear of pigeons (Nimpf *et al*., 2019; Nordmann *et al*., 2025). In principle, such a mechanism could be used by mole-rats if they possessed the required electrosensory molecular machinery, which has yet to be tested. Determining whether mole-rats rely on this mechanism or on other alternatives will be crucial for elucidating the sensory basis of mammalian magnetoreception.

## MATERIAL & METHODS

### Samples

For all experiments, eyes of sacrificed animals (Tab. S2) were dissected after perfusion. Potential iron contaminations of the tools and vials were minimized by immersion in 10% hydrochloric acid (HCl) over night and subsequent washing in ultrapure MilliQ water. For all experiments the magnetotactic bacteria (MTB) *Paramagnetospirillum magnetotacticum* strain MS1 (Koziaeva et al. 2023; DZMZ, DSM3856), served as positive control. MTBs were extracted by placing a magnet at the side of a culture tube overnight and suspending the formed pellet in MilliQ water or fixative. The solution was centrifuged (6000 rcf, 20 min, 18°C), the supernatant removed and the pellet resuspended in MilliQ three times before suspending the final pellet in MilliQ water.

### Prussian blue staining

#### Experiment 1 (standard Prussian Blue protocol)

To repeat the standard Prussian blue protocol used in previous studies, mole-rat eyes (n = 10 animals, 12 eyes) were preserved in 4% PFA for 2 hours at room temperature (RT) and subsequently washed and stored in phosphate buffered saline (PBS, pH = 7.2) at 4°C. We used both eyes (L = left, R = right) of two animals (2654, 1641) and one eye each from additional eight animals (1334R, 7022L, 4261L, 6931R, 2372R, 2910R, 3517R, 1169L). In four eyes of four animals, which were sectioned *in toto*, the dorsal part of the cornea was marked with a small cut to allow for later alignment. Furthermore, three additional eyes of three animals were dissected and the corneas sectioned completely. The samples were dehydrated (90 min each in EtOH 70%, 80%, 95%, 99.8%, 3 × 10 min xylene), infiltrated (2 × 120 min paraffin) and embedded in paraffin (Surgipath Paraplast 56°C, Carl Roth) using a tissue processor (STP120, Thermo Scientific) and an embedding station (Sakura Tissue-Tek TEC). Eyes were sectioned with a thickness of 10 µm at a rotary microtome (HM340, Microm) using ceramic coated blades (BLM00103P, DuraEdge) and mounted on silanized glass slides. Sections were deparaffinized for 3×10 min in xylene and rehydrated in a graded ethanol series (99.8%, 96%, 70%) and distilled water for 5 min per step, before staining for 2 × 30 min in Prussian blue staining solution (2% Hexacyanoferrate-II (Sigma-Aldrich, P9387) and 1% HCl in A. dest). Afterwards, sections were washed in Aq. dest. for 5 min and mounted in aqueous mounting media (RotiMount Aqua, Carl Roth). Some sections were counterstained with nuclear fast red for 5 min. Sections were dehydrated in a graded ethanol series for 5 min per step (30%, 50%, 70%, 96%, 99.8%, 100%), followed by Xylene two times for 10 minutes. Coverslips were placed on the slides with DPX new (Non-aqueous DPX mounting medium, Sigma-Aldrich, 1.00569.0500). All staining steps were performed at RT. The sections were manually screened for PB-positive particles at a transmitted light microscope (B201, Olympus) using 65x and 100x oil-immersion objectives. For each detected particle, we measured the distance from both edges of the section as well as the tissue localization.

#### Experiment 2 (enhanced Prussian blue protocol)

Mole-rat eyes (n = 3 animals) were preserved in 4% PFA for 2 hours at RT and subsequently washed and stored in phosphate buffered saline (PBS) at 4°C. The samples were dehydrated and embedded in paraffin (Paraplast, Sigma-Aldrich, P3558) as described above, using a tissue processor (TP1020, Leica) and an embedding station (Sakura Tissue-Tek TEC). Eyes were sectioned with a thickness of 5 µm at a rotary microtome (HM 355 S, Microm) using ceramic coated blades (BLM00103P, DuraEdge), and mounted on glass slides (Series I, Advances Adhesive, Trajan). Sections were deparaffinized for 5 min in Neo-Clear and rehydrated in a graded ethanol series (99.8%, 96%, 70%, 50%, 30%) and distilled water for 5 min per step, before transferring the sections into PBS for 10 min. For staining, slides were incubated in Prussian blue staining solution (2% Hexacyanoferrat-II ((Potassium hexacyanoferrate(II) trihydrate, Sigma-Aldrich, P9387)) and 1% HCl (pH=1) in A. dest) for 30 min and washed in 1x PBS three times for 5 min. To block endogenous peroxidases, slides were incubated with 0.3% H_2_O_2_ in methanol for 30 min. Staining was enhanced with 3,3,5,5 tetramethylbenzidine (TMB Substrate Kit, Peroxidase (HRP), Vector Laboratories, SK-4400) dripped onto the slides and incubated for 60 min. Subsequently slides were rinsed in PBS for 2 min and in A. dest for 10 min. Counterstaining was performed with nuclear fast red for 5 min. Sections were washed in PBS three times for 5 min and dehydrated in a graded ethanol series for 5 min per step (30%, 50%, 70%, 96%, 99.8%, 100%), followed by Xylene two times for 10 min. Slides were coverslipped in DPX new (Non-aqueous DPX mounting medium, Sigma-Aldrich, 1.00569.0500). To verify the methods on MTBs, samples were prepared as outlined above, dripped on microscopic slides and air dried. For MTBs staining followed the same protocol, but the deparaffinization and counterstaining was omitted. Furthermore, for some MTB samples enhancement was done with 3,3’-diaminobenzidine (DAB Substrate Kit, Peroxidase (With Nickel), Vector Laboratories, SK-4100) instead of TMB. All staining steps were performed at RT. Sections were digitized at 1000x magnification and a resolution of 55 nm/pixel with an Olympus VS200 slide scanner equipped with an automated oil dispenser, and screened manually for Prussian-blue positive particles in OlyVIA 2.1.1 (Olympus). Particles that were not in the same focal plane as the tissue were excluded.

#### Synchrotron X-ray fluorescence microscopy (XFM)

For XFM four eyes were preserved in 2.5% glutaraldehyde and 2% PFA in PB buffer overnight at 4°C and subsequently washed in 0.1 M PB. Samples were dehydrated in an ethanol series (15 min each in 30%, 50%, 70%, 90%, 2 × 20 min in 100%) and propylene oxide (2 × 10 min), and transferred via a 1:1 mixture of propylene oxide and middle epon (6 h) into pure epon. Pure epon was changed after 1 night and a second time after 1 hour, before transferring the eyes into embedding forms with a hardened bottom layer of epon. Eyes were covered by epon and hardened in a heating cabinet for 48 hours at 70°C. The samples were trimmed and sectioned (2 µm thickness) with a Leica EM UC6 ultra microtome with diamond knives (DiATOME trim 90 trim knife and a DiATOME Histo Jumbo 6.0 mm knife). Sections were collected on Si-N membranes with a thickness of 1 µm (12204137, SiRN-7.5-525-3.0-1000, Silson Ltd.). One cornea (ID: FA8) was sectioned in coronal plane completely with sections mounted every 10 µm (n = 24 sections). One right (ID: FA2, n = 6) and two left eyes (ID: FA3, n = 5; FA5, n = 5) were sectioned half along the sagittal plane and up to six sections from each were collected on Si-N membranes at a distance of 50 µm each. MTBs prepared as outlined above and suspended in MilliQ, were directly applied to the Si-N membrane and dried overnight. To obtain quantitative elemental density maps, XFM was performed at the Australian Synchrotron (ANSTO) XFM-beamline (Howard *et al*., 2020) (Melbourne, Australia, beamtime: 18094) and at the Deutsche-Elektronen-Synchrotron (DESY) beamline P06 at PETRA III (Hamburg, Germany, beamtime: I-20211267). High-resolution XFM maps were obtained at the French National Synchrotron Facility (SOLEIL) Nanoscopium beamline (Paris, France, beamtime: 20210899). Measurements were taken with an energy of 10 keV (ANSTO, DESY, SOLEIL) above the X-ray absorption K edge of iron (Fe), which is 7.112 keV (https://www.ruppweb.org/Xray/elements.html). The step size ranged between 1–2 µm for measurements of whole sections and 0.5–0.1 µm for measurements of specific regions of interest (ROIs). The DESY experiments were conducted at PETRA III, a 6 GeV synchrotron radiation source, specifically at the hard X-ray microprobe undulator beamline P06. More information on the set-up is given in Falkenberg et al. (Falkenberg *et al*., 2017). The incident X-ray energy was 10 keV. Prefocusing compound refractive lenses (CRLs) and Kirkpatrick-Baez (KB) mirrors were used to focus the beam down to 3.0 µm × 780 nm (horizontal × vertical) with a flux of 3 × 10^11^ photons s^−1^. For x-ray fluorescence (XRF) detection, a Vortex ME4 in 45° geometry with Xspress 3 pulse processors was applied. The P06 XRF measurements were processed using the non-linear least-squares fitting algorithm implemented in PyMCA (Solé *et al*., 2007). A thin film multi-element reference material (AXO RF17-14-18c10) which contains elements with XRF emission lines covering the relevant experimental energy range was measured to calculate elemental calibration. The XRF signal of elements not directly included in the reference material was calibrated from the reference measurement using elemental parameters obtained from xraylib (Schoonjans *et al*., 2011). This resulted in calibration factors for each element which correlate the measured counts directly to areal densities (g/cm^2^). Areal density maps of measurements at DESY were exported in g/cm^2^, and at ANSTO in ng/cm^2^, while measurements at SOLEIL were not quantified. The tissues of interest were segmented based on Sulfur(S)-density–maps (as these allowed best differentiation of tissues) and the Fe-density maps were masked accordingly with the SegmentEditor (Pinter, Lasso and Fichtinger, 2019) in Slicer 5.61 (Fedorov *et al*., 2012). Mean, standard deviation (sd), minimum and maximum Fe-density within each segmented structure were extracted with the Slicer module Segment statistics. Potential single domain magnetite in the masked Fe-density-maps of each tissue were identified in Image J 1.54 (Schindelin *et al*., 2012) using the analyze particle tool. Thresholds corresponded to the Fe content expected for 20 magnetite crystals of 50 nm diameter (with an expected Fe content of 4.658 × 10^-7^ ng per crystal) present in a single pixel. This threshold varied between data from different facilities and beamtimes due to different pixel-sizes (DESY: 414.06 ng/cm^2^; ANSTO 2021: 931.6 ng/cm^2^; ANSTO 2023: 232.9 ng/cm^2^). The theoretical calculations were confirmed by XFM-measurements of MTBs, showing particles with the Fe-content expected for magnetosome chains (Fig. 1C). To estimate the detection limit and the background Fe-density, the maximal Fe-density was measured for an area of 100 × 100 pixels within the epon section outside of the tissue. The maximum Fe-densities in the background were 37 ng/cm^2^ (31 ± 5 ng/cm^2^, n = 6 sections) for DESY, 194 ng/cm^2^ (143 ± 29 ng/cm^2^, n = 7 sections) for ANSTO2021, and 39 ng/cm^2^ (18 ± 10 ng/cm^2^, n = 6 sections) for ANSTO2023, corresponding to the Fe-density expected respectively in the presence of 1.79, 4.17 and 3.31 magnetite crystals per pixel. The background values were considered in the interpretation and discussion of the XFM results, but not subtracted from the measured values, because of relatively high fluctuations of the maxima across samples (see mean ± SD) and in general mean values per section around zero (supplement). In the same 100×100 pixels background area the mean and standard deviation of Titanium (Ti) and Chromium (Cr) density was measured. Fe-particle overlapping with Ti and/or Cr levels > mean + 2 SD in the respective sections were interpreted as originating from contamination.

#### Quantum diamond microscopy (QDM)

Quantum diamond microscopy was used for high-resolution mapping of the magnetic susceptibility in ROIs. Therefore, the second half of the eye FA5 (used for XFM) was sectioned in sagittal plane with a thickness of 500 nm. The sections were transferred to a diamond sample containing nitrogen vacancy (NV) defects for magnetic imaging. QDM was performed at the University of Melbourne.

The diamond magnetic sensors were engineered by implanting nitrogen into thin (< 100 μm) Type IIa diamond substrates ([N] < 5 ppb), sourced from Delaware Diamond Knives. The first imaging sensor (QD1) was overgrown with a 2 μm isotropically pure ^12^C layer prior to implantation. The second sensor (QD3) was implanted into the received substrate. The ^14^N implantation energy and dose for QD1 and QD3 was 5 keV (5×10^12^ cm^-2^) and 4 keV (1×10^13^ cm^-2^), respectively. The implants were performed into the <100> diamonds at an angle of 7° to avoid nitrogen channeling in the diamond. Following the implantation the diamond samples QD1 and QD3 were annealed for 4 hours at 1200 °C and 1000 °C respectively. The ⟨100⟩diamond sensor was then mounted onto a custom printed circuit board with a microwave antenna patterned onto the glass coverslip for imaging. The diamond sample used to image the MTB was sourced from Applied Diamond and implanted and annealed under similar conditions to QD3.

For quantum magnetic imaging, we employed a custom inverted widefield microscope equipped with a 532 nm excitation laser and bespoke microwave (MW) delivery. The MWs were used to drive ground state spin transitions in the NV defects within the diamond. The MW signal was delivered using a broadband microwave omega shaped antenna fabricated on the glass coverslip by evaporating 10/100 nm of chrome/gold onto a thin glass coverslip. The diamond sensor is then glued above the antenna with UV (Norland Optical Adhesive 63). The optical excitation and spin state preparation was performed by a 532 nm laser (2 W, Opus 532, Laser Quantum) focused onto the back of the aperture of a 40x oil objective (1.3 NA, Plan Fluor, Nikon). The excitation field of view (FOV) was 50 µm × 50 µm with an optical power density of ∼ 3.5 MW cm^−2^. The red photoluminescence (PL) of the NVs was separated from the green excitation light by passing back through the objective and a dichroic filter (Semrock, FF605-Di02-25×36). Red light was further filtered with a 660 − 735 nm bandpass filter before imaging with a scientific CMOS camera (Andor Zyla). A permanent magnet was used for the applied magnetic field at a strength of ∼4 mT, and the measurements were performed at room temperature and in ambient atmosphere.

In total five FOVs were imaged for the cornea on QD1 and 20 FOVs were imaged for the cornea on QD3.

The optically detected magnetic resonance (ODMR) spectra are collected by applying a pulsed-ODMR sequence and collecting the fluorescence with a camera exposure time of 30 ms. Each point in the ODMR spectrum is probed with a 10 µs optical pulse to polarise the NVs into the |0⟩sublevel. The fluorescence intensity from the NV centres is monitored while sweeping the MW frequency across a range which can drive the NV |0⟩to | − 1⟩and | + 1⟩ground state transitions. The MW π-pulses were tuned to efficiently flip the NV spins between the |0⟩to | ± 1⟩states. A second reference measurement is performed and subtracted without the MW pulse as a form of common mode rejection. A custom Python program was used to extract the ODMR spectrum at each imaging pixel. The spectra were fit using a set of Lorentzian functions of the form,

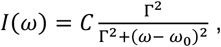

Where C is the contrast of the peak, Γ characterises the width of the Lorentzian and ω_0_ is the resonance frequency. Experimentally, maximal sensitivity is found by choosing the MW power that maximises the ratio of C to Γ (Dréau, 2011).

Each imaging pixel records an ODMR spectrum containing two ODMR transitions from the |0⟩to | − 1⟩and | + 1⟩states. The stray magnetic field 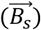 can be extracted from the separation between the transitions and knowledge of the applied magnetic field 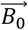. The total magnetic field, 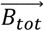, is measured directly via the Zeeman splitting of the | ± 1⟩states which can be calculated from the difference in the peak positions of the two Lorentzian peaks: 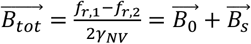. where *γ*_*NV*_ = 28.035 GHz/T is the gyromagnetic ratio of the NV centre, 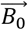 is the applied field (∼30 mT) and 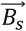 is the stray field of the sample projected along the NV axis. To account for artefacts in the magnetic field reconstruction due to variations in laser intensity across the FOV, 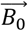 and MW inhomogeneity, a background correction is performed using a quadratic surface described in Broadway et al. 2020 (Broadway *et al*., 2020).

The component of the magnetic signal projected along the NV axis can be calculated from the peak-to peak value from a line scan through the stray magnetic field image. The peak-to-peak value can be found by fitting a Lorentzian, see below, the line scan.

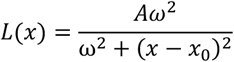

For dipole orientations that result in complex clover shaped projection, the combination of opposing peaks with the largest signal is chosen. For weak magnetic signals, fluctuations around zero can pose challenges to the fitting procedure, so a linear function, *f(x)*, is included in the fit. The total fit expression is therefore,

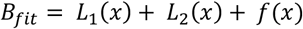

To find the peak-to-peak signal, we look to the minimum (H_2_) and maximum (H_1_) values of the fit function that occur at *x*_*2*_ and *x*_*1*_, respectively. This is not simply the amplitudes of the individual Lorentzians due to their overlapped regions and the influence of the linear part of the expression.

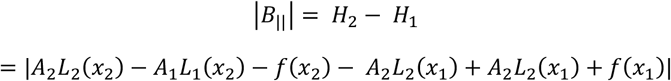

We can deduce the error in the peak-to-peak value from the fit standard errors, where the contribution to error from the widths is considered negligible.

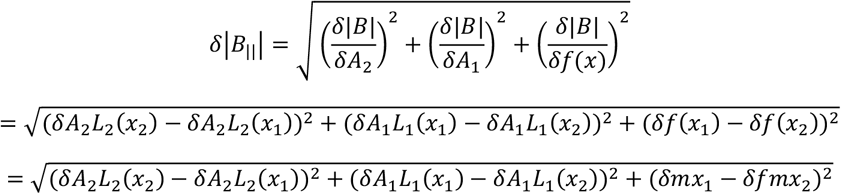

#### Energy-filtered transmission electron microscopy (EFTEM)

For screening via TEM and elemental analyses via energy filtered TEM (EFTEM) two eyes (Fa0251-L, FA0156-L) were fixed in 2.5% PFA/2% GA for 5 hours, and one eye (Fa0251-L) was post-fixed in 1% OsO_4_ for 2 hours, before embedding to increase contrast. 80 nm sections were cut with a diamond knife and collected on slot grids. To test the method, MTBs were prepared in the same way as for XFM and QDM (see above), dropped on a TEM grid with a magnet underneath for alignment of MTBs, and dried over night before imaging. TEM and EFTEM was performed at a Jeol JEM-2200F transmission electron microscope (JEOL GmbH) with x8k magnification. Magnification was chosen to resolve single domain magnetite crystals in MTBs while enabling a wide FOV (each 6 × 6 µm) for efficient screening. For two animals with two sections each, 91 FOVs were screened.

For elemental mapping, the jump-ratio for iron (Fe) was calculated, taking a post-edge image (width = 30 eV) after and one pre-edge image (width = 20 eV) before the electron absorption edge (708 eV for Fe (see EELS.info)). Parameters were optimized for the detection of magnetite crystals in MTBs. The following settings were used: Fe-post-edge: energy-shift = 725 eV, slit-width = 30 eV, exposure = 40 s; Fe-pre-edge: energy shift = 695 eV, slit-with = 20eV, exposure = 40 s. For the Fe-jump ratio the pre-edge image was divided by the post-edge image in Fiji/ImageJ. Furthermore, TEM images and zero-loss images were obtained for each ROI.

#### MRI-quantitative susceptibility mapping (QSM)

The magnetic susceptibilities of eyes (n = 8 eyes from different animals) and MTBs (n = 2) were mapped in 3D using MRI-QSM at the University of Graz. All samples were perfused with heparinized PBS (10 U/ml), fixed in 4% PFA in neutrally buffered 0.1 M PB for 1-2 h and subsequently washed and stored in PBS with 0.02% sodium azide to prevent bacterial growth. The eyes were placed in 3 mm NMR-tubes filled with PBS. MTB suspensions were prepared as outlined above, mixed 1:1 with agarose (2%) and filled into Hematocrit tubes (Micro-Hematocrit Capillary Tube, DWK KIMBLE). Measurements were performed on a 7 Tesla BioSpec MRI/MRS system (Bruker). A high-power gradient insert was employed, providing a maximum gradient strength of 660 mT/m and a slew rate of 4500 T/m/s. The RF system consisted of a cryogenically cooled transmit/receive RF coil to achieve high signal-to-noise ratio.

Quantitative susceptibility mapping data were acquired using a three-dimensional gradient-echo (3D GRE) sequence with the following parameters: repetition time = 60 ms, echo time = 6.5 ms, flip angle (α) = 12°, number of excitations = 24, acquisition matrix = 600 × 500 × 200, resulting in an isotropic spatial resolution of 20 µm × 20 µm × 20 µm, and receiver bandwidth = 82 kHz. QSM reconstruction was performed using a total generalized variation (TGV)-based algorithm as described by Langkammer and Bredies et al. (Langkammer *et al*., 2015) which enables robust susceptibility estimation via regularized inversion of the magnetic field-to-susceptibility dipole kernel.

For quantitative analysis of the MRI-QSM data the tissues of interest (cornea, retina, lens, ciliary body, n = 7, one eye (FA1) was excluded due to artifacts) as well as PBS background in three areas were manually segmented (Fig. 5B) in Slicer 5.6.1 (Fedorov *et al*., 2012) with the module Segment Editor (Pinter, Lasso and Fichtinger, 2019). Maximum, minimum, mean and median susceptibility, as well as the standard deviation were determined for each segmented structure with the module Segment Statistics for analysis in Python (V. 3.9). Based on this data the mean susceptibilities for each tissue over the 7 individuals was plotted and analyzed in Python (V. 3.9). Since the residuals met normality (Shapiro-Wilk: p = 0.965) and multiple tissues from the same individuals were measured, repeated measure-ANOVA was performed with Greenhouse-Geisser (ε = 0.485) correction for sphericity. As this revealed significant differences across tissues (p = 1.01×10^-7^), post-hoc pairwise paired t-tests were performed with Bonferroni correction to account for multiple testing (supplementary script1).

For visualization histogram data with voxel counts per susceptibility bin (here 1ppb) were exported for each segmented structure with a custom Python script (supplementary script2) run in the Python console of Slicer. An average histogram (binned to 10 ppb) for each tissue over the seven individuals and for the background, as well as scatterplots for each tissue, were plotted in Python (supplementary script3).

## DATA AVAILABILITY

Raw data (Density maps (XFM), MRI-QSM data), segmentations, the results of the quantification of the data, and the scripts used for the analysis will be deposited on EDMOND upon publication.

## ACKNOWLEDGEMENT

We are indebted to the core facilities at MPINB, in particular the electron microscopy facility (EMA) for assistance with sample preparation and data acquisition and all members of the Malkemper lab for helpful discussions. We thank Julia Herold for histological data acquisition. Part of this research was undertaken on the XFM beamline at the Australian Synchrotron, part of ANSTO (beamtime 18094). We acknowledge DESY (Hamburg, Germany), a member of the Helmholtz Association HGF, for the provision of experimental facilities. Parts of this research were carried out at PETRA III. Data was collected using beamline P06 operated/provided by DESY Photon Science. We would like to thank the P06 beamline staff for assistance during the experiments. Beamtime was allocated for proposal I-20211267, as well as a part of the P06 in-house research. This research was supported in part through the Maxwell computational resources operated at Deutsches Elektronen-Synchrotron DESY, Hamburg, Germany. We thank SOLEIL (beamtime 20210899) for beamtime and support. This project was funded by the Max Planck Society and the European Research Council (ERC StG No 948728 to EPM). E.P.W. and D.A.S. acknowledge that this research was supported by the Australian Research Council Centre of Excellence in Quantum Biotechnology through project number CE230100021 a. E.P.W. acknowledges that this research was supported by the Commonwealth through an Australian Government Research Training Program Scholarship [DOI: https://doi.org/10.82133/C42F-K220] and the Jean Laby Women in Physics Travel Award. D.A.S. would like to acknowledge support from the ARC Mid-Career Industry Future Fellowship (IM240100073) and the MCN Technology Fellow Ambassador Program. The Department of Mathematics and Scientific Computing of the University of Graz, with which KB is affiliated, is a member of NAWI Graz (https://www.nawigraz.at/en/).

## AUTHOR CONTRIBUTIONS

Conceptualization and design: EPM, LM, KN

Methodology: LM, KN, EPW, EPM, GF, DB, CD, KB, MB, DH, DJP, SI, DS

Investigation: LM, KN, EPW, AS, EB, AL, GF, DB, CD, KB, MB, DH, DJP, KM, SWK, LZ, MMMD, EPM

Formal analysis: LM, KN, EPW, AS, EB, AL, GF, DB, CD, KB, MB, DH, DJP, KM, SWK, LZ, MMMD, EPM

Resources: DS, SB, SI,

Data Curation: LM, KN

Writing – Original draft: LM, EPM

Writing – Review & Editing: all authors

Visualization: LM, EPW

Supervision: EPM

Funding acquisition: EPM, EPW, DAS, LM (beamtime proposals), KN (beamtime proposals)

