## Supplemental Information for "Multimodal imaging reveals no evidence for magnetite-based magnetoreceptors in the mole-rat eye"

- **Supplementary Text**
- **Supplementary Figures S1 to S7**
- **Supplementary Tables S1 to S2**
- **Supplementary References**

### Supplementary Text

#### Can MRI microscopy detect magnetite nanocrystal compounds in mole-rat eyes?

**Introduction.** Mole-rats are suspected to sense the magnetic field using magnetite nanocrystals. Are these nanocrystals detectable using MRI? We answer this question in two steps: First, we estimate the magnetic moment of the nanocrystals, and then we assess their impact on the MRI microscopy signal following (1).

**Magnetic moment of nanocrystal compound.** First, we estimate the magnetic moment of a compound of 20 nanocrystals. These crystals have a mass of 0.012875 pg. The saturation magnetization of magnetite is 90 A m $kg^{-1}$ (2). Thus, the magnetic moment of the 20 nanocrystals turns out to be

$$\mu = 90 \text{ A m kg}^{-1} \cdot 0.012875 \text{ pg} = 1.16 \text{ mA}\mu\text{m}^2$$

Note that this is approximately a factor 100 smaller than the magnetic moment of dopaminergic neurons at 9.4 T.

**MR detectability of a nanocrystal compound.** Here, we assess whether a compound of 20 nanocrystals is detectable with MRI microscopy. The experimental setup considered is a 7 T MRI system, which acquires a single echo gradient echo image at echo time 6.5 ms at an isotropic resolution of 20 µm. We approximated the crystal compound as a cubic magnetization inclusion of 1 $\mu m^3$ volume in the center of the imaging volume, adjusting the magnetization such that the cube had the same magnetic moment as estimated in Eq. 1. Note that all calculations were performed in the static dephasing approximation, so the estimated impact of the nanocrystal compounds is an upper bound of the effect. We simulated the complex-valued gradient echo signal using a numerical simulation soon to be published. Figure S7 shows the resulting magnitude and phase images. The dephasing of the nanocrystal compound reduces the MRI signal by 37 % in the central voxel, but only by less than 3 % in the neighboring voxels. In the voxels directly neighboring the central voxel, the phase difference was in the range 0.08 rad to 0.16 rad. In QSM maps at the MRI imaging resolution and magnetic field strength, the nanocrystal compound would appear as a susceptibility inclusion of 26 ppb. The estimated effect of the nanocrystal compounds should be compared with the recorded MRI microscopy data to assess whether it is detectable. For detectability, the impact of the nanocrystal compounds should be larger than the noise in the images, but also larger than the variability of the MRI signal due to other parts of the mole-rat eye.
Based on the calculation a single chain of 20 sd crystals would appear as a susceptibility inclusion of 26 ppb (1 sdp = 1.3 ppb). This lays within the fluctuations of the PBS background susceptibility, ranging from -199.8 – 89.7 ppb (mean -17.2 ± 44.5 ppb, n = 103875 voxels). Therefore, the detection limit lays at 71.1 ppb (mean + 2 SD of PBS background, corresponds to 55 sdp). In conclusion, while MRI-QSM provides a qualitative overview of the susceptibility of different tissues within the mole-rat eye, it would only reliably detect magnetoreceptors if they contain a larger number of single-domain crystals in a voxel.

**Supplementary Figures**

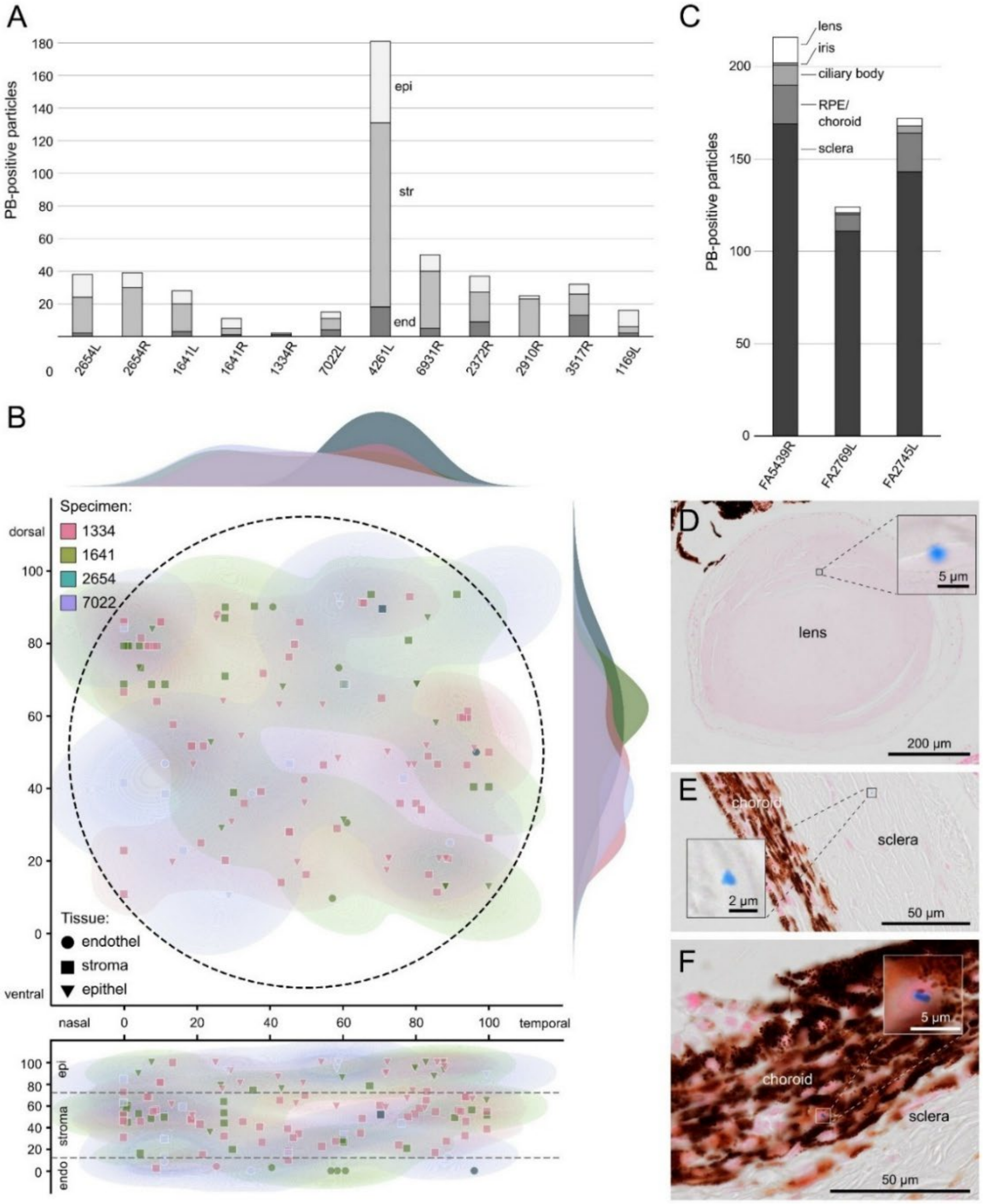


**Fig. S1. Prussian blue (PB) screening of the mole-rat eye reveals iron particles with a random distribution. A & B** Screen of the cornea using the classical PB protocol. **A** Quantification of PB-positive particles in the corneas of ten animals, highlighting large inter-individual differences. n = 612 sections (2654L = 56; 2654R = 69; 1641L =58; 1641R = 17; 1334R = 17; 7022L = 75; 4261L = 49; 6931R = 77; 2372R = 55; 2910R = 32; 3517R = 47; 1169L = 60). **B** Distribution of PB-positive particles within eight corneas of four animals showed no regular pattern regarding layers or position (n = 970 sections; 1334L = 45; 1334R = 72; 1641L = 170; 1641R = 128; 2654L = 137; 2654R = 165; 7022L = 173; 7022R = 80). The x- and y axes show the relative position from nasal and ventral (top) or basal (bottom) C–F Screening of other ocular tissues using a TMB-enhanced PB protocol. **C** Quantification of PB-positive particles in the different tissues of complete eyes of three animals (n = 960 sections; FA5439R = 365; FA2769L = 291; FA2745L = 304). **D–F** Exemplary particles in the (D) lens, (E) sclera, and (F) choroid.

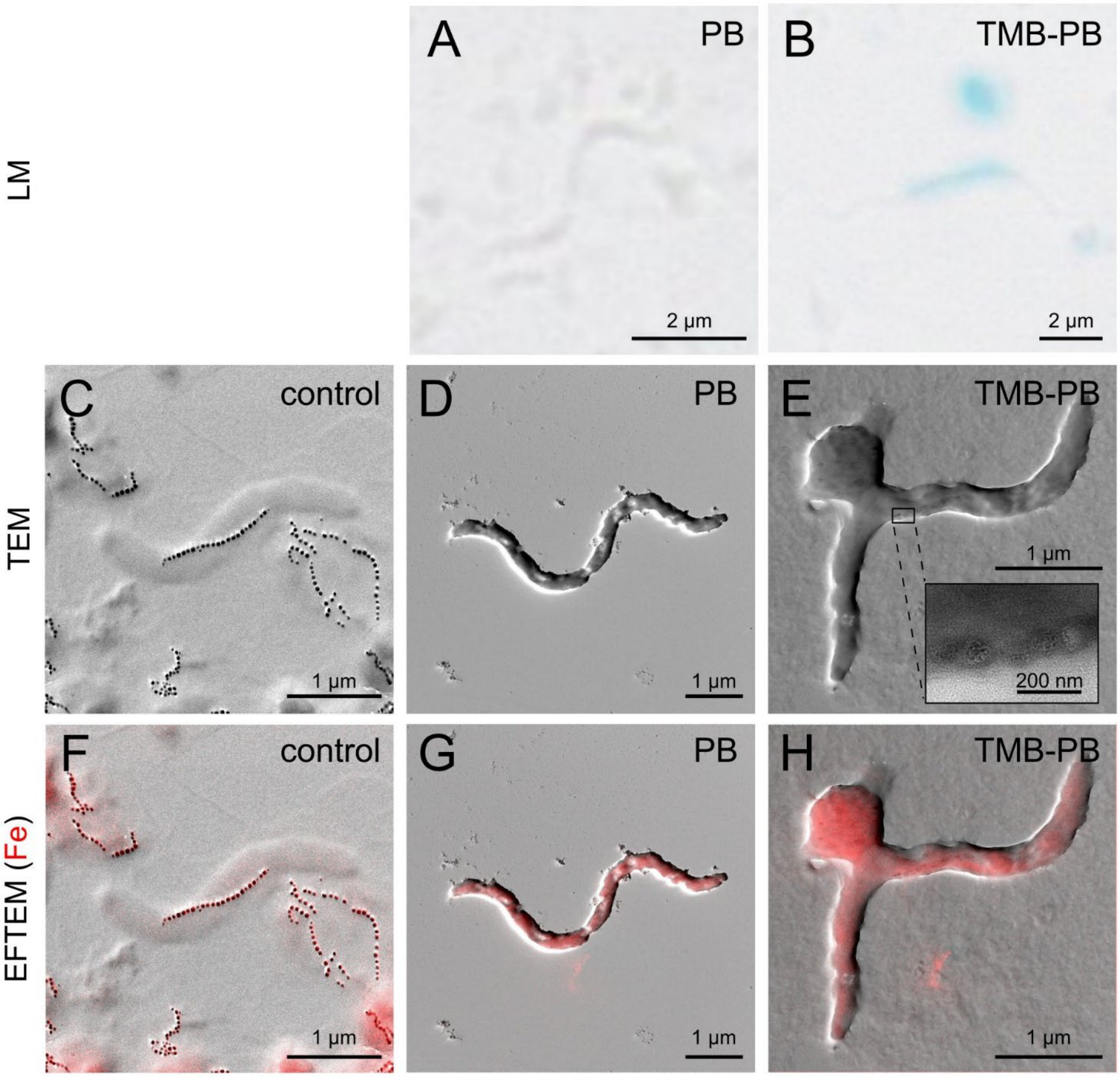


**Fig. S2. Magnetotactic bacteria (*Paramagnetospirillum magnetotacticum*) stained with different versions of the Prussian blue method. A–C** Light microscopy (LM) images of MTBs treated with the classical (A) and the TMB-enhanced (B) Prussian-blue (PB) protocols show that amplification is necessary to visualize magnetite in MTBs. **C–E** Transmission electron microscopy (TEM) images of untreated (C), PB-stained (D) and PB-TMB stained (E) MTBs show that magnetite crystals partially dissolve after treatment. F–H Fe-jump ratio (EFTEM (Fe)) of untreated MTBs (F) and MTBs treated with the classical PB protocol (G) and the TMB enhanced Prussian blue protocol (H). Prussian blue treatment dissolved the magnetite within the MTBs, leading to a homogeneous distribution of Fe (in red) within the cells, rather than concentrations in the magnetosomes, as indicated by the Fe-jump-ratio. The Fe-jump ratio shows the qualitative distribution of iron and is not quantitative. Note that some MTBs have turned ovoid in culture and do not show the typical spirillum form but still possess magnetosomes.

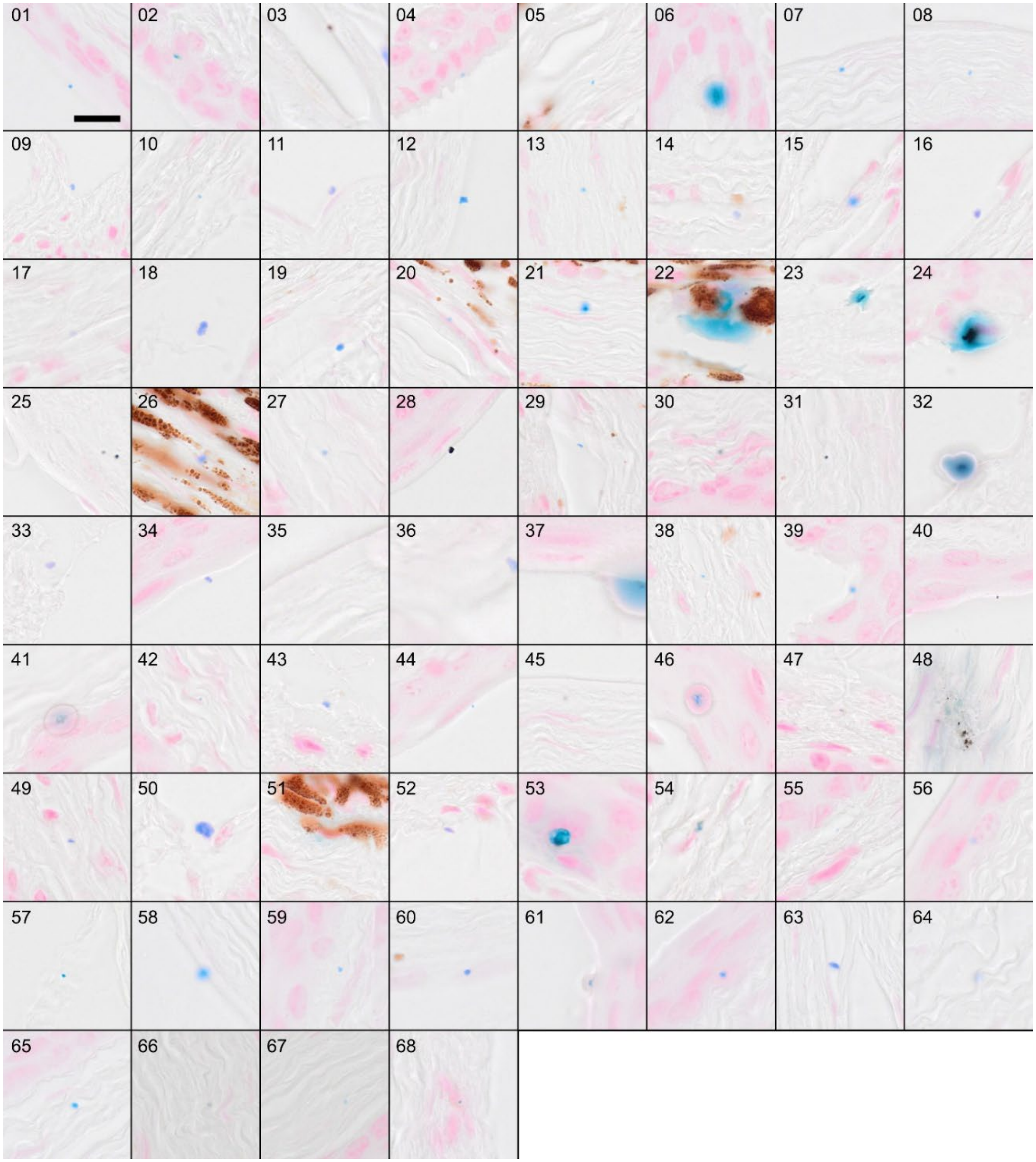


**Fig. S3. All 68 Prussian blue (PB) positive particles in the cornea of three mole-rat eyes, found with the TMB-enhanced staining, show variable shapes, sizes, and localizations.** Same samples as shown in Fig. 2 and Fig. S1C-F; 01-26 = FA5439R; 25-52 = FA2769L; 53-68 = FA2745L; All particles are shown at the same magnification in a 27.5 x 27.5 µm window. Scale (in 01) = 10 µm.

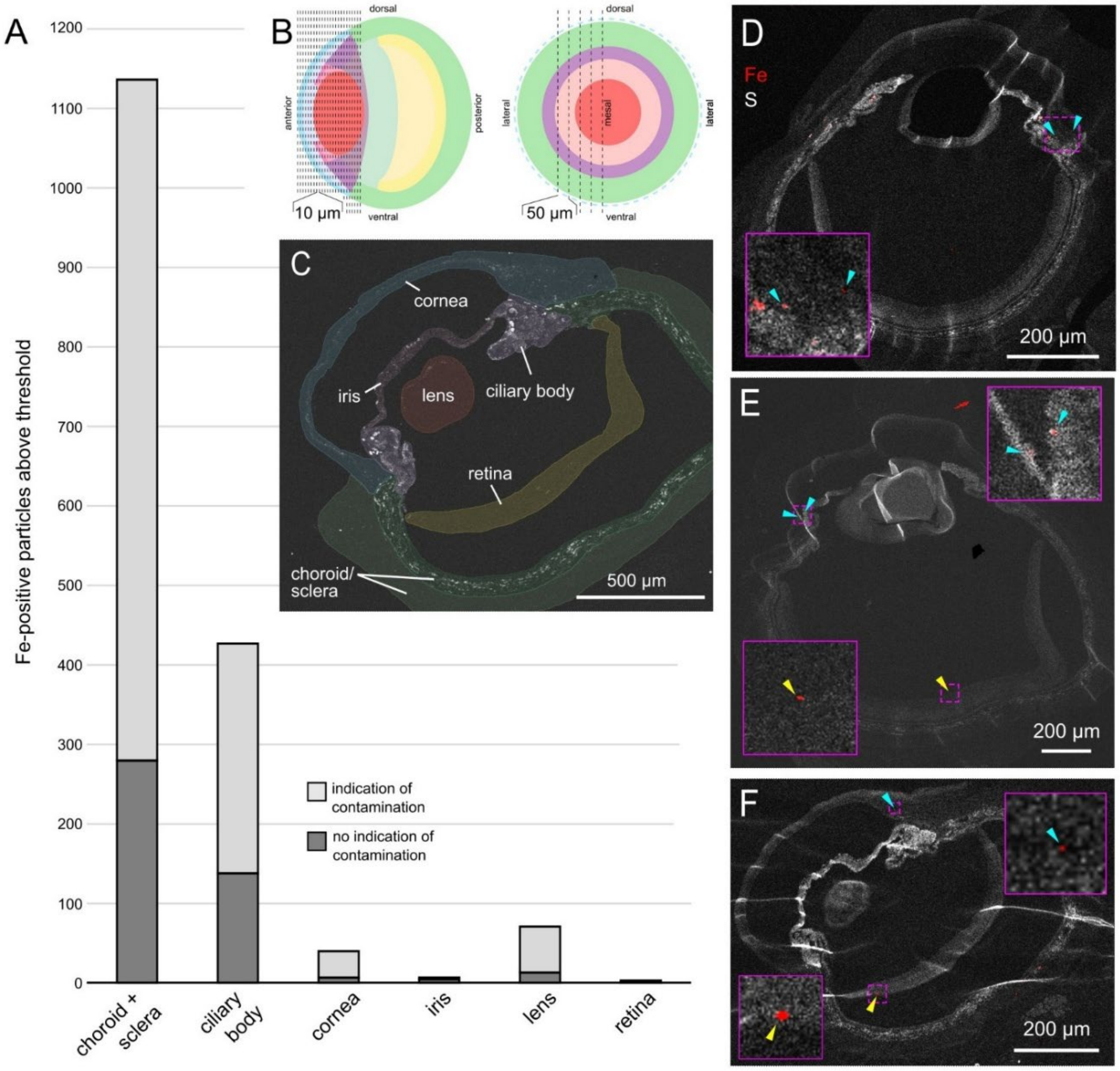


**Fig. S4.** Elemental density maps based on synchrotron X-ray fluorescence microscopy (XFM) of 2 µm epon sections of mole-rat (Fukomys anselli) eyes on Si-N membranes. **A** Particles with Fe-density above that expected for 20 sd crystals/pixel, with and without signs of contamination. n = total number of sections from four animals (n = 136 sections; choroid + sclera = 35; ciliary body = 35; cornea = 19; iris = 11; lens = 24; retina = 14). **B** Schematic representation of the eye in sagittal (left) and coronal (right) view with coronal (left) and sagittal (right) sections indicated by dashed lines; colors follow C. **C** Fe-density map with analyzed areas marked and colorized. **D–F** S-density maps with exemplary particles found above threshold without signs of contamination in the cornea (blue arrows) and retina (yellow arrows) measured at ANSTO. These particles are mainly located at the boundary of the cornea and ciliary body (D & E), on folds (E), or on top of (E) or underneath (F) the retina.

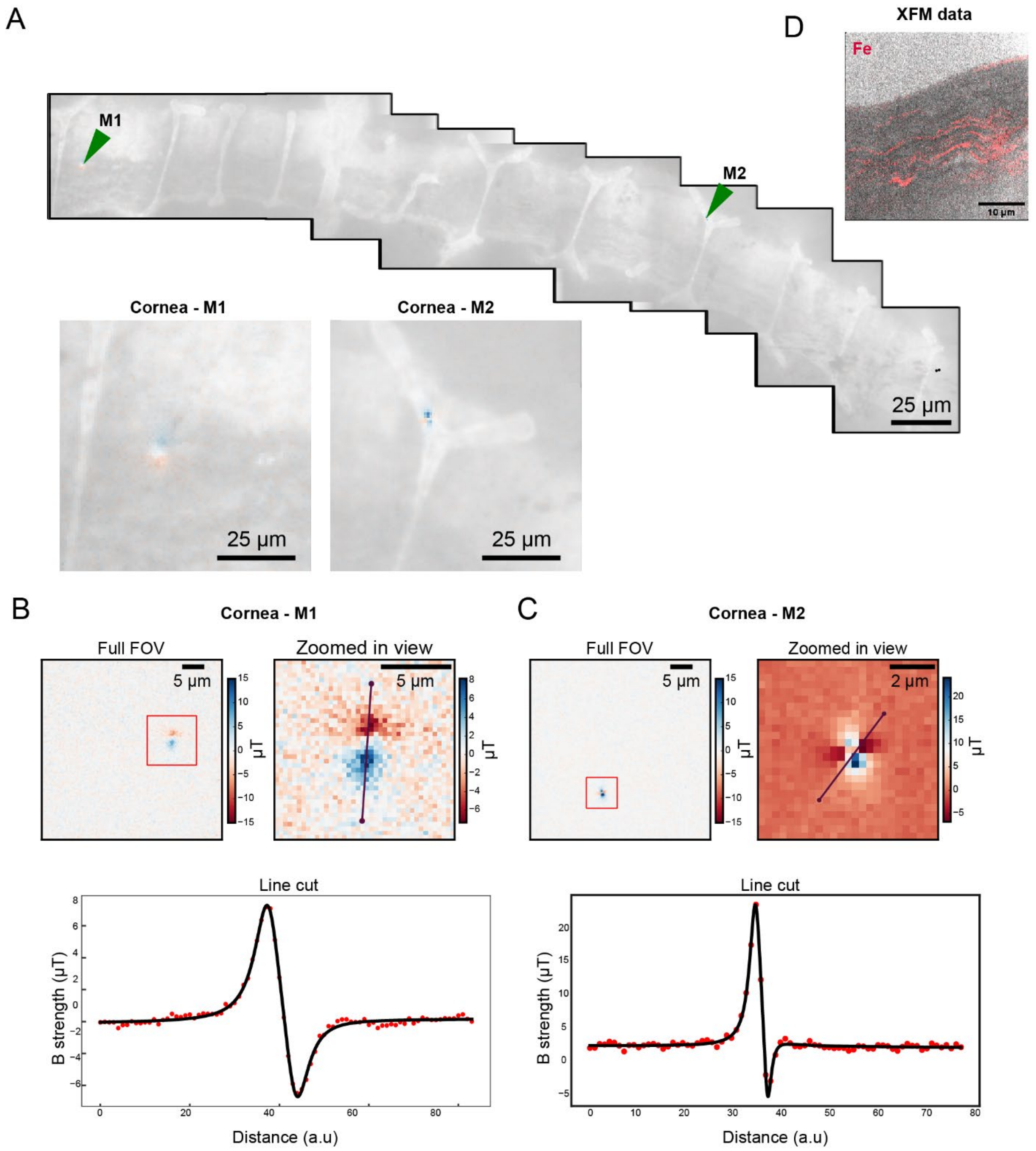


**Fig. S5. Quantum diamond imaging of a mole-rat cornea section. A** Fluorescent image showing the location of the ferrimagnetic signal (indicated by green arrows) in the quantum diamond measurement of the cornea. Note that M1 is clearly visible by light microscopy and M2 colocalizes with a fold in the section. **B** and **C** Stray magnetic field maps for the magnetic particles (M1, M2) with line cut through the particles. **D** Note the absence of ferrimagnetic signals in the corneal stroma where iron rich lines were detected with XFM.

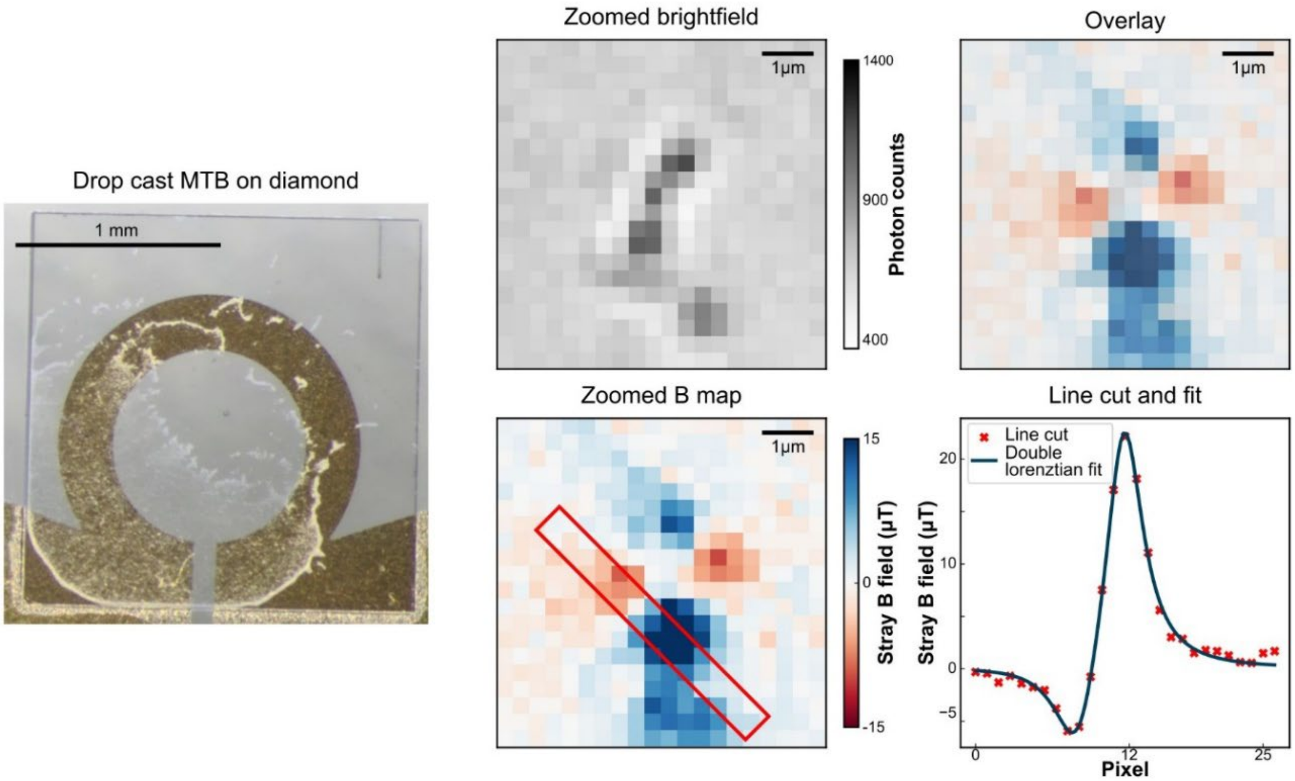


**Fig. S6. Stray magnetic field maps of magnetotactic bacteria (*Paramagnetospirillum magnetotacticum*) drop-casted onto a diamond sample.** The red box in the zoomed B map indicates the line cut.

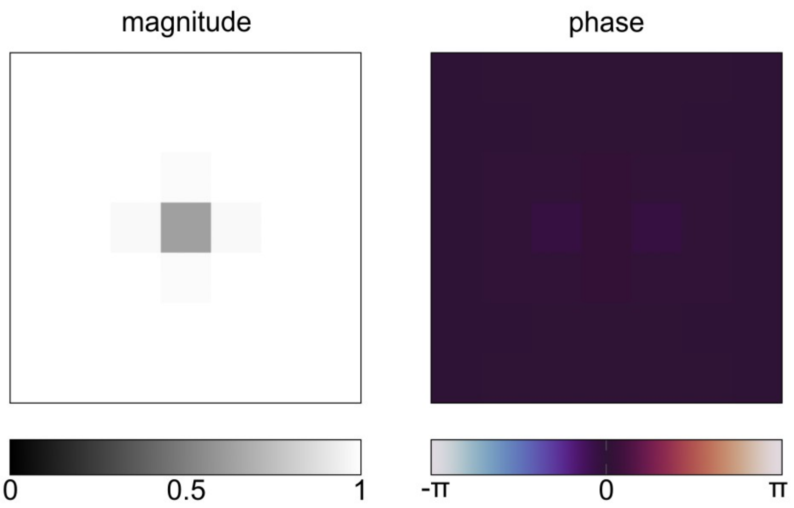


**Fig. S7. Magnitude and phase of the simulated MRI signal around a compound of 20 nanocrystals.**

### Supplementary Tables

**Table S1.** Average Fe-density (ng/cm$^2$) in different ocular tissues based on XFM-measurements. n indicates the number of individuals.

| Tissue | FA2 | FA3 | FA5 | FA8 | Average | SD | n (animals) |
|---|---|---|---|---|---|---|---|
| **Ciliary body** | 35.53 | 31.21 | 30.90 | 54.50 | 38.04 | 11.18 | 4 |
| **Iris** | 36.01 | 16.78 | 18.54 | 54.72 | 31.51 | 17.74 | 4 |
| **Choroid** | 22.10 | 21.09 | 18.49 | 18.64 | 20.30 | 2.16 | 4 |
| **Sclera** | 3.47 | 3.69 | 1.84 | 8.73 | 4.43 | 2.98 | 4 |
| **Cornea** | 2.12 | 4.11 | 2.92 | 7.91 | 4.26 | 2.56 | 4 |
| **Lens** | na | 0.74 | -0.11 | 7.49 | 2.71 | 4.17 | 3 |
| **Retina** | 1.67 | 0.43 | 1.24 | na | 1.11 | 0.63 | 3 |

**Table S2.** Material studied. Abbreviations: F = female, L = left, M = male, R = Right.

| Animal ID | Sex | Eye | Sample name | Method | Tissue | Sampling | Experimenter | Thickness |
|---|---|---|---|---|---|---|---|---|
| 5365 | M | L | FA8 | MRI-QSM | Eye | Complete (3D) | C.D., K.N. | |
| 5366 | M | R | FA7 | MRI-QSM | Eye | Complete (3D) | C.D., K.N. | |
| 8470 | M | L | FA2 | MRI-QSM | Eye | Complete (3D) | C.D., K.N. | |
| 30439 | F | R | FA3 | MRI-QSM | Eye | Complete (3D) | C.D., K.N. | |
| 30471 | M | l | FA6 | MRI-QSM | Eye | Complete (3D) | C.D., K.N. | |
| 30475 | M | R | FA5 | MRI-QSM | Eye | Complete (3D) | C.D., K.N. | |
| 35356 | F | L | FA4 | MRI-QSM | Eye | Complete (3D) | C.D., K.N. | |
| 353562 | M | R | FA1 | MRI-QSM | Eye | Complete (3D) | C.D., K.N. | |
| 1334 | F | R | 1334R | Prussian-blue (standard) | Eye | 17 sections | A.S. | 10 µm |
| 1641 | M | L | 1641L | Prussian-blue (standard) | Eye | 58 sections | A.S. | 10 µm |
| 1641 | M | R | 1641R | Prussian-blue (standard) | Eye | 17 sections | A.S. | 10 µm |
| 2654 | F | L | 2654L | Prussian-blue (standard) | Eye | 56 sections | A.S. | 10 µm |
| 2654 | F | R | 2654R | Prussian-blue (standard) | Eye | 69 sections | A.S. | 10 µm |
| 7022 | M | L | 7022L | Prussian-blue (standard) | Eye | 75 sections | A.S. | 10 µm |
| 4261 | M | L | Fa14 4261L | Prussian-blue (standard) | Cornea | 49 sections | J.H. | 10 µm |
| 6931 | M | R | FA23R | Prussian-blue (standard) | Cornea | 77 sections | J.H. | 10 µm |
| 2472 | F | R | FA34R | Prussian-blue (standard) | Cornea | 55 sections | J.H. | 10 µm |
| 1169 | M | L | FA34 1169L | Prussian-blue (standard) | Eye | 60 sections | J.H. | 10 µm |
| 3517 | F | R | FA34 3517R | Prussian-blue (standard) | Eye | 47 sections | J.H. | 10 µm |
| 2190 | M | R | FKA14R | Prussian-blue (standard) | Eye | 32 sections | J.H. | 10 µm |
| 2745 | M | L | FA2745L (2021_28) | Prussian-blue (TMB) | Eye | Complete (304 sections) | E.B., A.L. | 5 µm |
| 2769 | M | L | FA2769L (2021_26) | Prussian-blue (TMB) | Eye | Complete (291 sections) | E.B., A.L. | 5 µm |
| 5439 | M | R | FA5439R (2021_25) | Prussian-blue (TMB) | Eye | Complete (365 sections) | E.B., A.L. | 5 µm |
| 5366 | M | L | FA7 | XFM (ANSTO) | Cornea & retina | Wholemount | K.N. | 2 µm |
| 8470 | M | R | FA2 | XFM (ANSTO) | Eye | 6 sections | K.N. | 2 µm |
| 30439 | F | L | FA3 | XFM (ANSTO) | Eye | 5 sections | K.N. | 2 µm |
| 30471 | M | R | FA6 | XFM (ANSTO) | Cornea & retina | Wholemount | K.N. | 2 µm |
| 30475 | M | L | FA5 | XFM (ANSTO) | Eye | 5 sections | K.N. | 2 µm |
| 35356 | F | R | FA4 | XFM (ANSTO) | Cornea & retina | Wholemount | K.N. | 2 µm |
| 353562 | M | L | FA1 | XFM (ANSTO) | Cornea & retina | Wholemount | K.N. | 2 µm |
| 5365 | M | R | FA8 | XFM (DESY) | Eye | 24 sections | K.N. | 2 µm |
| 251 | F | L | QD1 | QDM | Eye | 1 section | E.W., L.M. | 500 nm |
| 251 | F | L | QD2 | QDM | Eye | 1 section | E.W., L.M. | 500 nm |
| 251 | F | L | QD3 | QDM | Eye | 1 section | E.W., L.M. | 500 nm |
